# Transgenic Mouse Models Establish a Protective Role of Type 1 IFN Response in SARS-CoV-2 infection-related Immunopathology

**DOI:** 10.1101/2022.12.17.520843

**Authors:** Nishant Ranjan Chauhan, Soumya Kundu, Ramyasingh Bal, Diya Chattopadhyay, Subhash Mehto, Rinku Sahu, Rina Yadav, Sivaram Krishna, Kautilya Kumar Jena, Sameekshya Satapathy, Krushna C Murmu, Bharati Singh, Saroj Kumar Das, Sarita Jena, Krishnan H Harshan, Gulam Hussain Syed, Punit Prasad, Santosh Chauhan

## Abstract

Type 1 interferon (IFN-I) response is the first line of host defense against invading viruses. In the absence of definite mouse models, the role of IFN-I in SARS-CoV-2 infections remained to be perplexing. Here, we developed two mouse models, one with constitutively high IFN-I response (hACE2; *Irgm1^−/−^*) and the other with dampened IFN-I response (hACE2; *Ifnar1^−/−^*) to comprehend the role of IFN-I response during SARS-CoV-2 invasion. We found that hACE2; *Irgm1^−/−^* mice were resistant to lethal SARS-CoV-2 infection with substantially reduced cytokine storm and immunopathology. In striking contrast, a severe SARS-CoV-2 infection along with immune cells infiltration, inflammatory response, and enhanced pathology was observed in the lungs of hACE2; *Ifnar1^−/−^* mice. Additionally, hACE2; *Ifnar1^−/−^* mice were highly susceptible to SARS-CoV-2 neuroinvasion in the brain accompanied by immune cell infiltration, microglia/astrocytes activation, cytokine response, and demyelination of neurons. The hACE2; *Irgm1^−/−^ Ifnar1^−/−^* double knockout mice or hACE2; *Irgm1^−/−^* mice treated with STING or RIPK2 pharmacological inhibitors displayed loss of the protective phenotypes observed in hACE2; *Irgm1^−/−^* mice suggesting that heightened IFN-I response accounts for the observed immunity. Taken together, we explicitly demonstrate that IFN-I protects from lethal SARS-CoV-2 infection, and *Irgm1* (IRGM) could be an excellent therapeutic target.

**GRAPHICAL ABSTRACT:** 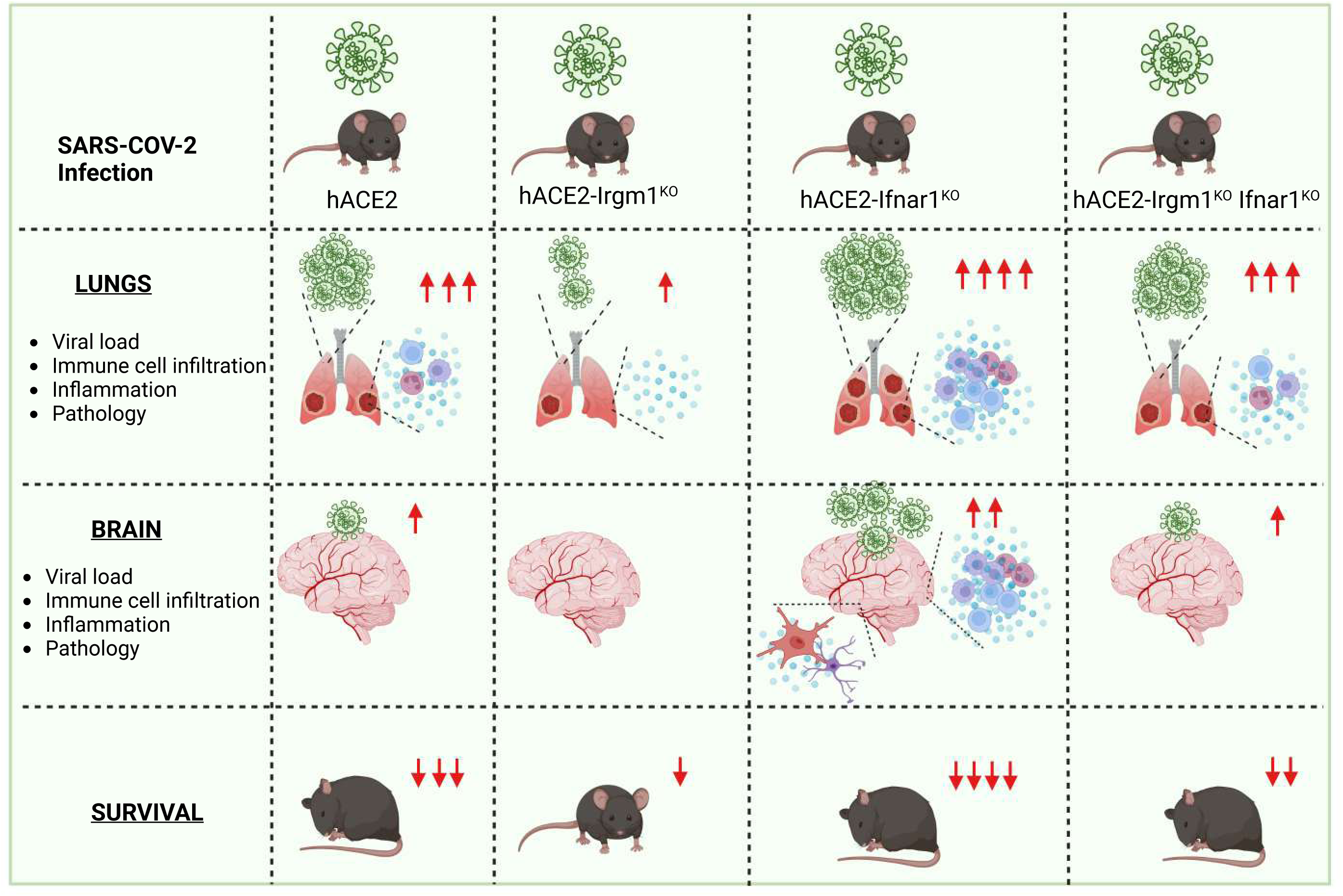

## Introduction

Severe acute respiratory syndrome coronavirus 2 (SARS-CoV-2) is an enveloped, positive-strand RNA virus that caused the Coronavirus disease 2019 (COVID-19) pandemic and killed several million worldwide (Dong et al, 2020). The clinical manifestation of the disease is variable, ranging from asymptomatic infection to severe lung infection and multi-organ failure to death (Hu et al, 2021). Understanding the basis of this heterogeneity in disease outcomes could be enormously important to save the vulnerable population from SARS-CoV-2 variants and other similar deadly viruses in the future.

Type I interferons are the most potent first lines of innate immune defense against invading viruses. Recent population-wise studies suggest that genetic defects leading to deficiency (loss-of-function) of Type I interferons (IFN-I) receptors (IFNAR1/2) in human populations increases susceptibility to viral infections including SARS-CoV-2 and results in poor clinical outcomes (Abolhassani et al, 2022; Bastard et al, 2022; Duncan et al, 2022; Khanmohammadi et al, 2022; Meyts, 2022; Zhang et al, 2022). Also, IFN-I-neutralizing autoantibodies were detected in ∼10% of patients with severe COVID-19 pneumonia but not in individuals with asymptomatic infections (Bastard et al, 2021; Bastard et al, 2020; Frasca et al, 2022). These clinical studies indicate that enhancing the IFN-I response could be an important prophylactic measure to restrict SARS-CoV-2 infection. Thus, genetically modified mouse models need to be developed for testing this hypothesis and understanding the mechanisms.

Using different *in vitro* and *in vivo* models, both protective and pathogenic functions of IFN-I were proposed in COVID-19 disease (Anjum et al, 2021; Boudewijns et al, 2020; Domizio et al, 2022; Israelow et al, 2020; King & Sprent, 2021; Mao et al, 2022). Using a mouse model based on adeno-associated virus (AAV)–mediated expression of human angiotensin I-converting enzyme 2 (hACE2), Israelow et al. concluded that IFN-I does not control SARS-CoV-2 replication but drives pathological responses (Israelow et al, 2020). Similarly, several early COVID-19 studies found that IFN-I is induced during the SARS-CoV2 infection or in patient samples, and based on this correlation, it was suggested that IFN-I response may aggravate inflammation and pathology in COVID-19 (Chiale et al, 2022; Kim & Shin, 2021; Lee et al, 2020). On the other hand, later studies suggested the protective function of the IFN-I response including the genetic evidence pointed out above. A RIG-I agonist by inducing IFN-I response was shown to protect from acute and chronic SARS-CoV-2 infection in the hACE2 mouse model (Chiale et al, 2022; Mao et al, 2022). Although several studies have been conducted, in the absence of accurate animal models, the role of interferons in COVID-19 disease remains unclear.

SARS-CoV-2 infection is triggered by the binding of the spike protein to hACE2, triggering lung injury, inflammation, and subsequent respiratory distress (Yan et al, 2020; Ziegler et al, 2020). Even though several animal species (Hamsters, ferrets, and non-human primates) have been tested for SARS-CoV-2 susceptibility in laboratory studies, none of these exhibited the severe disease symptoms seen in hospitalized patients (Cleary et al, 2020; Jia et al, 2021; Rockx et al, 2020). The parental SARS-CoV-2 strain or variants do not infect standard laboratory mice due to differences in ACE2 receptors between mice and humans (Letko et al, 2020; Wan et al, 2020). Transgenic mice expressing the hACE2 receptor under the control of the K18 gene promoter (K18-hACE2) exhibit symptoms that are most closely related to those experienced by humans (Arce & Costoya, 2021; Golden et al, 2020; Oladunni et al, 2020; Winkler et al, 2020; Yinda et al, 2021). Hence, it was suggested to be the most suitable model to study mechanisms of pathogenesis or screening of prophylactic and therapeutic remedies (Arce & Costoya, 2021; Golden et al, 2020; Oladunni et al, 2020; Winkler et al, 2020; Yinda et al, 2021).

COVID-19 is considered to be primarily a respiratory disease, however, a large number of patient and animal studies suggest that SARS-CoV-2 can invade the central nervous system to trigger neuropathology and cognitive impairments (Bauer et al, 2022; Douaud et al, 2022; Jacob et al, 2020; Meinhardt et al, 2021; Rutkai et al, 2022; Seehusen et al, 2022; Sepehrinezhad et al, 2021; Song et al, 2021; Veleri, 2022). SARS-CoV-2 could enter the nervous system by crossing through the neural-mucosal interface in the olfactory mucosa (Meinhardt et al, 2021), or passing through the blood-brain barrier (Krasemann et al, 2022; Rhea et al, 2021; Zhang et al, 2021). The factors and mechanisms contributing to SARS-CoV2 neuroinvasion remain unclear, and hence the development of new mouse models that could enhance our understanding of this process would be beneficial.

Immunity-related GTPase M 1 (Irgm1) is a master negative regulator of IFN-I response by regulating the cGAS-STING and RIG-I-MAVS pathways (Jena et al, 2020; Rai et al, 2021). Thus, Irgm1 depletion induces an antiviral IFN state in cells resulting in resistant to the infection with several viruses (Nath et al, 2021). Also, IRGM (human orthologue of Irgm1) depleted THP1 cells were found to be resistant to SARS-CoV-2 infection (Nath et al, 2021). However, it is unclear whether *Irgm1* knockout (*Irgm1^−/−^*) mice are protected from SARS-CoV-2 infection and subsequent immunopathology. It is imperative to gain this knowledge since it will not only shed light on the role that IFN-I plays in controlling SARS-CoV-2 infections but will also provide insight into whether Irgm1 is a viable therapeutic target against COVID-19.

To unequivocally define the role of IFN-I response in SARS-CoV-2 infection, we generated hACE2 expressing *Irgm1^−/−^* mice (hACE2; *Irgm1^−/−^*) with constitutively high IFN response and hACE2 expressing *Ifnar1^−/−^* mice (hACE2; *Ifnar1^−/−^*) with blunted IFN-I response. We found that hACE2; *Irgm1^−/−^* mice were highly protected from SARS-CoV-2 infection, cytokine storm, and subsequent immunopathology. In contrast, hACE2; *Ifnar1^−/−^* mice were highly susceptible to SARS-CoV-2 invasion in the lungs leading to increased cytokine storm, immune cell infiltration, and severe pathology causing rapid death. In the absence of IFN-I response in hACE2; *Ifnar1^−/−^* mice, SARS-CoV-2 could invade the brain resulting in immune cell infiltration, hyperactivation of microglial cells, hyperinflammation, and rapid demyelination of neurons. To delineate whether the protective phenotype observed in hACE2; *Irgm1^−/−^* is due to IFN-I response, we generated hACE2; *Irgm1^−/−^ Ifnar1^−/−^* double knockout mice, which we found to be highly susceptible to SARS-CoV-2 infection resulting in high mortality. Also, inhibition of IFN-I using STING and RIPK2 inhibitor in hACE2; *Irgm1^−/−^* mice as well as siRNA knockdown of anti-viral factors (ISG15, Oas1a, Viperin, etc) in *Irgm1^−/−^* macrophages enhanced SARS-CoV-2 replication suggesting their role in protecting *Irgm1^−/−^* cells/mice from viral invasion. Additionally, we show that Irgm1 depletion more effectively guards against SARS-CoV-2 infection as compared to prophylactic IFN-α treatment, and thus, *Irgm1* could be an excellent therapeutic target against SARS-CoV-2 infection. Taken together, these independent mouse models and cell-based studies demonstrate that an intact IFN-I response is essential to defend against SARS-CoV-2 infection, and an enhanced IFN-I response could be highly protective against invading SARS-CoV-2 and subsequent life-threatening respiratory and neurological manifestations.

## Results

### hACE2; Irgm1^−/−^ mice are protected from lethal SARS-CoV-2 infection and pathology

The *Irgm1^−/−^* mice (Collazo et al, 2001; Jena et al, 2020; Liu et al, 2013; Mehto et al, 2019) were crossed with the K18-hACE2 (JAX Strain #:034860; **hereafter, hACE2**) transgenic mice to generate hACE2 (hemizygous) expressing *Irgm1* knockout mice (**hereafter, hACE2*; Irgm1^−/−^***). Throughout this study, hemizygous hACE2 mice were used for the SARS-CoV-2 infection. The hACE2 and hACE2*; Irgm1^−/−^* were inoculated with 5X10^4^ (low-dose) or 5X10^5^ (high-dose) plaque-forming units (PFU) via the intranasal route with highly infectious early SARS-CoV-2 strain of clade 19A (Kumar et al, 2021; Singh et al, 2022; Suresh et al, 2021) (Genbank Accession MW559533). For the low-dose experiment, the mice were sacrificed on the 6 and 15 days post-infection (DPI), whereas for the high-dose experiment, mice were sacrificed on the 7 DPI (**Figure 1A**). As expected, the uninfected mice gained body weight during the course of infection, while the SARS-CoV-2-infected mice lost weight substantially (**Figure 1B-1D**). Compared to hACE2 mice, reduction of hACE2; *Irgm1^−/−^* mice weight was significantly less in both low-dose and high-dose experiments (**Figure 1B-1D**). In the low-dose 15 days experiment, a progressive weight loss of hACE2 mice was observed and several (5/10) mice died by the 8 DPI (**Figure 1D**). The rest of the surviving hACE2 mice except one started regaining weight (**Figure 1D**). Similar results were obtained previously with the hACE2 mouse model (Golden et al, 2020; Yinda et al, 2021).

**Figure 1.**
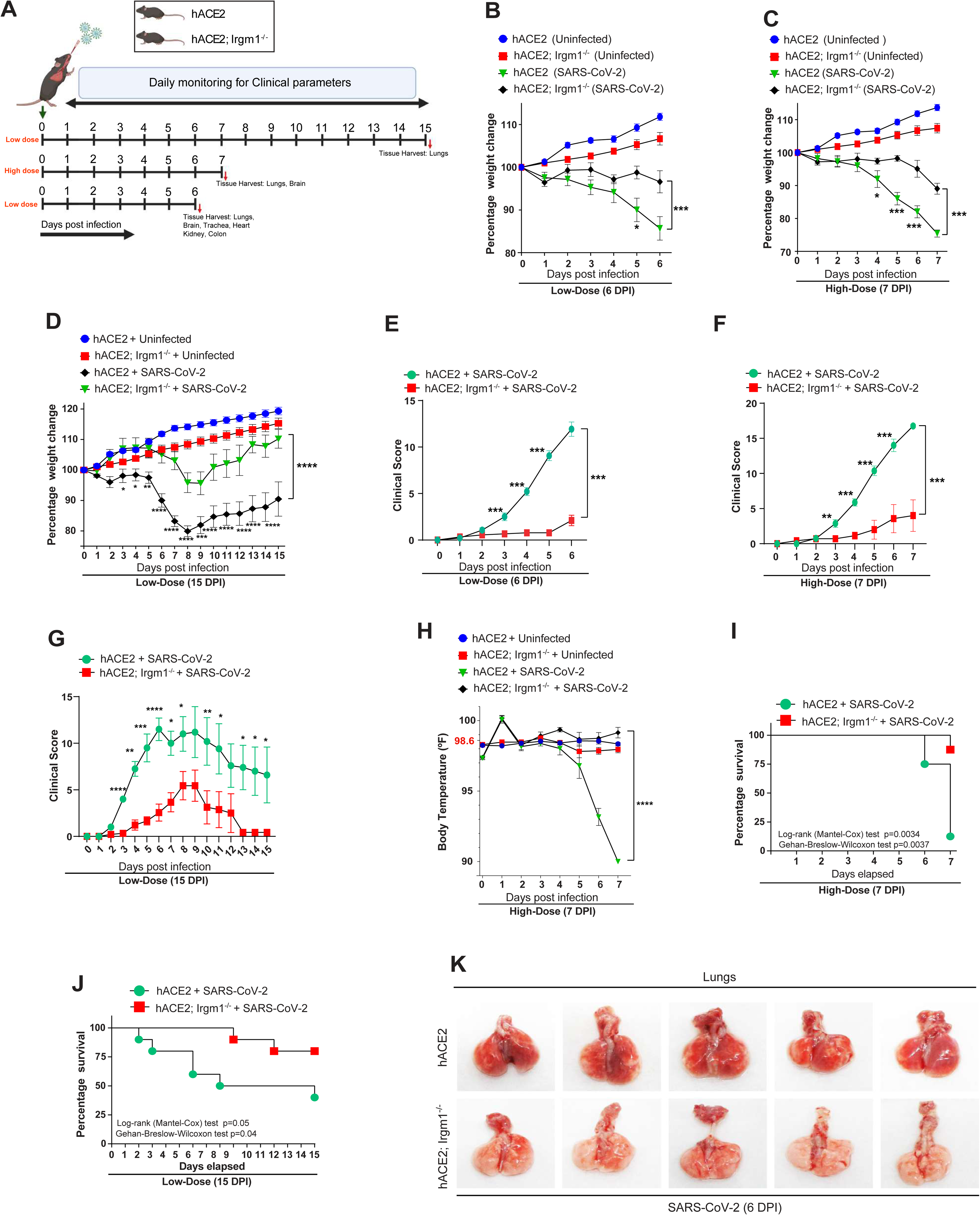
hACE2; Irgm1^−/−^ transgenic mice are protected from severe SARS-CoV-2 infection. **A.** Schematic representation of the experimental design depicting the timeline of SARS-CoV-2 infection in hACE2 and hACE2; *Irgm1*^−/−^ transgenic mice. Mice were intranasally inoculated with 5X10^4^ PFU (low dose) and 5X10^5^ PFU (high dose) of SARS-CoV-2. Image created using Biorender.com. **B-D.** Graph depicts percentage change in the body weight of uninfected (n=8) and SARS-CoV-2 infected hACE2 (n=8) and hACE2; *Irgm1^−/−^* (n=7) mice **(B)** Low dose for 6 days, **(C)** High dose for 7 days **(D)** Low dose for 15 days. Mean ± SEM, *p < 0.05, **p < 0.01, ***p < 0.001 and ****p < 0.0001, Two-way ANOVA (Tukey’s multiple comparison test). **E-G.** Graph showing clinical scores (**E)** of hACE2 (n=14) and hACE2; *Irgm1^−/−^* (n=9) mice infected with low dose of SARS-CoV-2 for 6 days, **(F)** of hACE2 (n=8) and hACE2; *Irgm1^−/−^* (n=7) mice infected with high dose of SARS-CoV-2 for 7 days or **(G)** of hACE2 (n=8) and hACE2; *Irgm1^−/−^* (n=9) mice infected with low dose of SARS-CoV-2 for 15 days. Mean ± SEM, *p < 0.05, **p < 0.01, ***p < 0.001 and ****p < 0.0001, Two-way ANOVA followed (Sidak’s multiple comparison test). **H.** Graph depicting core body temperature (in °F) of SARS-CoV-2 infected hACE2 (n=6) and hACE2; *Irgm1^−/−^* (n=6) mice. Mean ± SEM, *****p < 0.00001, Two-way ANOVA (Tukey’s multiple comparison test). **I, J.** Kaplan–Meier survival graph showing percentage survival of **(I)** hACE2 (n=8) and hACE2; *Irgm1^−/−^* (n=8) mice inoculated with a high dose of SARS-CoV-2 and monitored for 7 days [**p = 0.0034, Log-rank (Mantel-Cox) test, **p = 0.0037, Gehan-Breslow-Wilcoxon test], and **(J)** hACE2 (n=10) and hACE2; *Irgm1^−/−^* (n=10) mice inoculated with a low dose of SARS-CoV-2 and monitored for 15 days [*p = 0.05, Log-rank (Mantel-Cox) test, *p = 0.04, Gehan-Breslow-Wilcoxon test]. **K**. Lung images showing gross pathology in hACE2 and hACE2; *Irgm1^−/−^* mice infected with SARS-CoV-2 (5X10^4^ PFU/mice) at 6 dpi.

In hACE2 mice, the development of clinical symptoms associated with SARS-CoV-2 infection begins with body shivering, which is followed by hunched posture and labored breathing before they become completely unresponsive and need to be euthanized **(Supplementary Video 1**; **Supplementary Figure 1A and 1B**). In both low-dose (**Figure 1E**) and high-dose (**Figure 1F**) infection experiments, the majority of hACE2; *Irgm1^−/−^* mice exhibited very mild clinical symptoms (**Supplementary Video 2**). In the 15 DPI experiments, clinical scores reached a maximum on the 7^th^ DPI and then improved for the surviving mice (**Figure 1G**). While the clinical scores for all the surviving hACE2; *Irgm1^−/−^* mice were fully restored to normal by 13 DPI, the hACE2 mice still displayed abnormal clinical scores (**Figure 1G**). Hypothermia (a rapid drop in body temperature) was found to be associated with the increased mortality rate in COVID-19 patients (Fatteh et al, 2021; Maait et al, 2021). We observed a rapid drop in temperature in hACE2 mice, but not in hACE2; *Irgm1^−/−^* mice (**Figure 1H**). In the high-dose infection experiment, only one hACE2 mouse survived by 7DPI and in striking contrast only one hACE2*; Irgm1^−/−^* mouse died by 7 DPI (**Figure 1I**). In the low-dose infection experiment, 50% of mice succumbed to SARS-CoV-2 infection by 8 DPI, whereas no mortality was noticed in hACE2*; Irgm1^−/−^* mice by this day (**Figure 1J**). Thus, hACE2*; Irgm1^−/−^* exhibited higher survival percentage compared to hACE2 mice in both low and high-dose model.

A striking difference was observed in the gross lung pathology of mice sacrificed at 6 and 7 days post-infection, where lungs from hACE2 mice displayed extensive inflammatory (necrotic lesions) areas; lungs of hACE; *Irgm1^−/−^* mice were clear. (**Figure 1K and Supplementary Figure 1C**). The hACE2 mice that survived (4 out of 10) in 15 DPI experiments displayed less inflammation, indicating eventual clearance of viral infection **(Supplementary Figure 1D**). Overall, we found that *Irgm1^−/−^* mice were protected from debilitating and lethal SARS-CoV-2 challenges.

### hACE2; Irgm1^−/−^ mice exhibit reduced SARS-CoV-2 burden and attenuated immunopathology

We then evaluated SARS-CoV-2 RNA levels in the lungs of infected mice in four separate experiments that were terminated at 5 DPI, 6 DPI, 7 DPI, and 15 DPI. In all experiments, hACE2 mice lacking *Irgm1^−/−^* had significantly lower viral RNA levels than hACE2 mice (**Figure 2A-2D**). In agreement, in Western blotting experiments, where an excessive amount of SARS-CoV-2 nucleocapsid (NC) protein was detected in hACE2 mice lung lysates, little or no NC protein was detected in hACE2; *Irgm1^−/−^* mice (**Figure 2E-2H and Supplementary Figure 1E**). A similar result was obtained ‘with immunofluorescence in lung sections (**Figure 2I-2J and Supplementary Figure 1F**) demonstrating that hACE2; *Irgm1^−/−^* mice have a significantly lower viral load than their hACE2 mice. SARS-CoV-2 infection considerably induced Irgm1 protein levels in the lungs (**Figure 2E, 2G, and Supplementary Figure 1E**), which is consistent with our previous in vitro finding that viral infection induces *Irgm1* expression(Nath et al, 2021). In the 15 DPI experiment, a very low viral copy number was detected in surviving (4 out of 10) hACE2 mice lungs (**Figure 2D**), which corresponds with the reversal of clinical score (**Figure 1G**), weight loss (**Figure 1D**) and pathology (**Supplementary Figure 1D**) in these mice. These results demonstrate that hACE2; *Irgm1^−/−^* mice are resistant to SARS-CoV-2 invasion.

**Figure 2.**
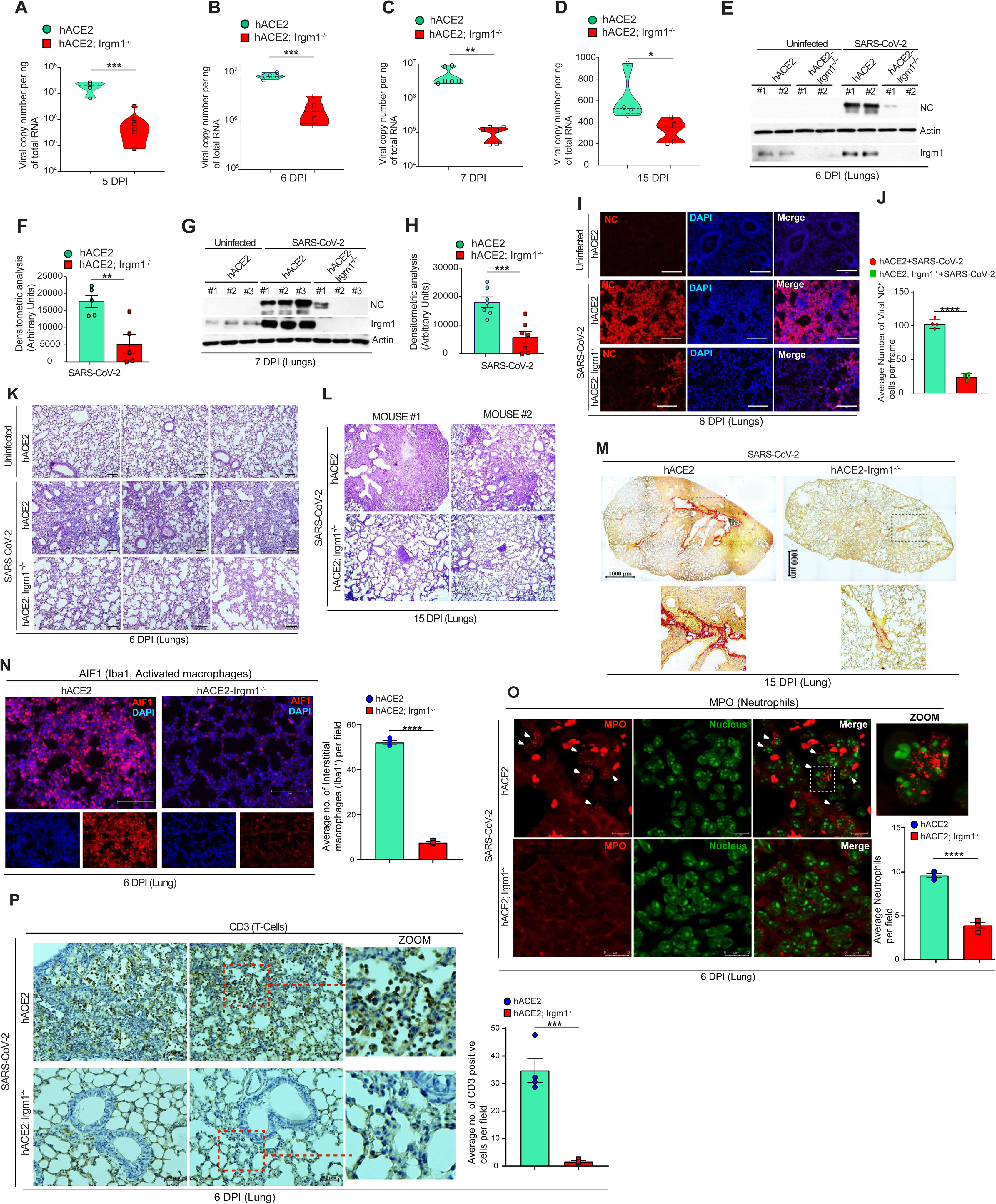
Reduced SARS-CoV-2 burden and mild immunopathology in hACE2; Irgm1^−/−^ mice. **A-D.** Viral copy number analysis from total RNA isolated from lungs of SARS-CoV-2 infected hACE2 and hACE2; *Irgm1^−/−^* mice at **(A)** 5 dpi (n=6), **(B)** 6 dpi (n=4), **(C)** 7 dpi (n=6), and **(D)** 15 dpi (n=4, 5). Mean ± SEM, *p < 0.05, **p < 0.01, ***p < 0.001, Student’s unpaired t-test. **E-H.** Western blot analysis of nucleocapsid (NC) protein expression in lung lysates of uninfected and SARS-CoV-2 infected hACE2 and hACE2; *Irgm1^−/−^* mice at **(E)** 6 dpi **(G)** 7 dpi. **(F, H)** Bar graphs depicting densitometric analysis (using ImageJ) of western blots normalized with beta-actin (mean ± SEM, **p < 0.01, ***p < 0.001, Student’s unpaired t-test). **I-J. (I)** Representative images of SARS-CoV-2 nucleocapsid (NC) immunostained lungs sections of uninfected hACE2 and SARS-CoV-2 infected hACE2 and hACE2; *Irgm1^−/−^* mice (6 dpi). **(J)** Bar graphs showing an average number of NC positive cells per field (10 fields from each experiment, n=4 experiments, mean ± SEM, ****p < 0.0001, Student’s unpaired t-test). **K-L.** Representative hematoxylin & eosin stained lung images of uninfected hACE2 and SARS-CoV-2 infected hACE2 and hACE2; *Irgm1^−/−^* mice at **(K)** 6 dpi and (**L)** 15 dpi. **M.** Representative picrosirius red stained lung images of SARS-CoV-2 infected hACE2 and hACE2; *Irgm1^−/−^* mice showing collagen deposition (in red) at 15 dpi. **N.** Representative images of AIF1 (IBA1, activated macrophage) immunostained lungs sections of SARS-CoV-2 infected hACE2 and hACE2; *Irgm1^−/−^* mice (6 dpi). Right panel shows bar graphs for average number of IBA1 positive cells per field (10 fields from each experiment, n=4 experiments, mean ± SEM, ****p < 0.0001, Student’s unpaired t-test). **O.** Representative confocal images of MPO (activated neutrophils) immunostained lungs sections of SARS-CoV-2 infected hACE2 and hACE2; *Irgm1^−/−^* mice (6 dpi). Right panel shows bar graphs for average number of MPO positive cells per field (10 fields from each experiment, n=4 experiments, mean ± SEM, ****p < 0.0001, Student’s unpaired t-test). **P.** Representative immunohistochemistry (IHC) images of CD3 (activated T-Cell) immunostained lungs sections of SARS-CoV-2 infected hACE2 and hACE2; Irgm1^−/−^ mice (6 dpi). Right panel shows bar graphs for average number of CD3 positive cells per field (10 fields from each experiment, n=4 experiments, mean ± SEM, ***p < 0.001, Student’s unpaired t-test).

To assess the degree of lung injury upon SARS-CoV-2 infection, we performed a histopathological evaluation using hematoxylin and eosin (H&E) staining. In the lungs of hACE2 mice, extensive lung damage with alveolar space consolidation, alveolar septa thickening, parenchymal collapse, and excessive infiltration of lymphoid cells was observed (**Figure 2K and Supplementary Figure 2A**). In hACE2; *Irgm1^−/−^* mice, the lung injury was strikingly less as compared to hACE2 mice (**Figure 2K and Supplementary Figure 2A**). In 15 DPI lungs, although viral load was low (**Figure 2D**), the injury was much more prominent in hACE2 mice with extensive consolidation, necrosis, and fibrosis (**Figure 2L and Supplementary Figure 2B**). Here again, the lung pathology was considerably less in hACE2; *Irgm1^−/−^* mice (**Figure 2L and Supplementary Figure 2B**).

Lung fibrosis status was evaluated using Picrosirius red (PSR) stain, which is commonly used for collagen staining. In the 6 DPI experiment, although severe lung consolidation was evident in hACE2 mice, no prominent collagen deposition was observed (**Supplementary Figure 2C**). Whereas, in 15 DPI lungs of hACE2 mice a large number of prominent fibrotic regions were evident in consolidated areas (**Figure 2M**). In contrast, a very mild PSR staining was observed in hACE2; *Irgm1^−/−^* mice lungs (**Figure 2M**). These results were confirmed using immunohistochemistry (IHC) staining with a specific collagen-I antibody (**Supplementary Figure 2D**). The results show that the lungs of hACE2; *Irgm1^−/−^* mice show significantly low fibrosis as compared to hACE2 mice.

Respiratory viruses, including SARS-CoV-2, induces excessive infiltration of immune cells in the lungs. Thus, we assessed immune cell infiltration in SARS-CoV-2-infected lungs of hACE2 and hACE2; *Irgm1^−/−^* mice using standard markers. The activated macrophages were stained using AIF1/Iba1 (Allograft inflammatory factor 1 or Ionized calcium-binding adapter molecule 1), neutrophils by MPO (Myeloperoxidase), and T-cells by CD3 in the lung sections (**Figure 2N-2P**). In contrast to hACE2; *Irgm1^−/−^* mice, hACE2 mice lungs showed significantly higher infiltration of macrophages (in alveolar, bronchial, interstitial, and vascular regions), neutrophils (mainly in bronchial, interstitial, and vascular regions), and T-cells (mainly in bronchial, interstitial, and vascular regions) (**Figure 2N-2P**). Taken together, the results show severe immunopathology and lung damage in hACE2 mice due to an acute inflammatory environment. In contrast, hACE2; *Irgm1^−/−^* mice exhibited significantly less immune cell infiltration, inflammation, pathology, and lung damage.

Various vital organs were also monitored for SARS-CoV-2 infection. A low to very low level of viral RNA was detected in the colon, heart, and kidney of mice (**Supplementary Figure 3A-3C**). Although hACE2; *Irgm1^−/−^* mice organs showed consistently lower infection than hACE2 mice, the difference was insignificant (**Supplementary Figure 3A-3C**). Tracheal viral loads were higher than other organs (**Supplementary Figure 3D-3E**). Like the lungs, the trachea of hACE2; *Irgm1^−/−^* mice displayed significantly reduced infection rates (**Supplementary Figure 3D-3E**). There was very high variability in viral loads in the brains of hACE2 mice. Some hACE2 mice brains showed very high SARS-CoV-2 RNA levels, while others had negligible levels (**Supplementary Figure 3F**). Although the viral loads in hACE2; *Irgm1^−/−^* mice brains were lower than in hACE2 mice, the difference was insignificant. To corroborate the finding, we performed Western blotting with brain lysates using NC antibody. Where none of the hACE2; *Irgm1^−/−^* mice brain was positive, NC band was detected in 4 out of 17 hACE2 mice brains (**Supplementary Figure 3G**). This data indicates that where there is a limited capacity of SARS-COV-2 to invade the brain of hACE2 mice, no SARS-CoV-2 neuroinvasion was detected in hACE2; *Irgm1^−/−^* mice brains.

Overall, these results demonstrate that *Irgm1^−/−^* mice are highly resistant to SARS-CoV-2 infection, exhibiting considerably lower immunopathology and lung damage.

### hACE2; Irgm1^−/−^ mice are less susceptible to SARS-CoV-2 delta variant infection

We then tested whether *Irgm1* knockout mice could also resist SARS-CoV-2 delta infection, a deadly variant associated with high mortality (Mlcochova et al, 2021). The mice were infected with 5X10^4^ PFU of the SARS-CoV-2 delta variant for 6 days. The hACE2; Irgm1*^−/−^* mice demonstrated much milder clinical symptoms (whole body shivering/shaking, hunched posture, abnormal labored breathing, unresponsive to touch) than hACE2 mice (**Supplementary Figure 4A**). Interestingly, where the weight of hACE2 mice dropped rapidly, the average weight of hACE2; *Irgm1^−/−^* mice was increased (**Supplementary Figure 4B**). The majority of hACE2 mice reached euthanasia conditions by 6 DPI, whereas none of the hACE2; *Irgm1^−/−^* mice did (data not shown). In agreement with clinical symptoms, hACE2 mice had substantial gross pathology in their lungs as compared to hACE2; *Irgm1^−/−^* mice (**Supplementary Figure 4C**). The lungs of the hACE2 mice were highly inflamed with several visible lesions (**Supplementary Figure 4C**). As detected by qRT-PCR and western blotting, hACE2; *Irgm1^−/−^* mice had significantly lower viral loads in lungs than hACE2 mice (**Supplementary Figure 4D and 4E**). A higher degree of alveolar space consolidation, thickening of the alveolar septa, parenchymal collapse, and excessive infiltration of immune cells was observed in hACE2 mice as compared to hACE2; *Irgm1^−/−^* mice (**Supplementary Figure 4F**). Together, our findings suggest that depletion of *Irgm1* protects even from the dreaded SARS-CoV-2 delta strain.

### hACE2; Irgm1^−/−^ mice exhibit mild cytokine responses to SARS-CoV-2 infection

Extensive cytokine response upon SARS-CoV-2 infection is a hallmark of COVID-19 disease leading to severe pathology and acute damage to the lungs. We performed RNA sequencing analysis to evaluate the transcriptional landscape of uninfected and SARS-CoV-2 infected (6 DPI and 7DPI) lungs of hACE2 and hACE2; *Irgm1^−/−^* mice (**Supplementary Figure 5A**). Principal component analysis (PCA) revealed distinct transcriptional signatures of all the groups (**Supplementary Figure 5B**). A striking difference between the transcriptome (overall) of the infected hACE2 and hACE2; *Irgm1^−/−^* lungs was observed (**Supplementary Figure 5B**).

First, we compared the transcriptional landscape of uninfected vs infected lungs of hACE2 mice. The differentially expressed genes (DEGs, p<0.05) that were induced more than 3-folds in infected lungs (compared to uninfected) (**Supplementary Dataset 1**) were subjected to gene ontology (GO) enrichment analysis using Metascape pathway analysis tool (Zhou et al, 2019) (**Figure 3A, left panel**) and Reactome pathway analysis tool (Jassal et al, 2020)(**Supplementary Figure 5C**). An extensive number of innate /adaptive immunity-related pathways including cytokine/chemokine response, interferon (α, β, and γ) response, anti-viral response, leukocytes activation/adhesion/migration, neutrophil degranulation, antigen processing/presentation were overrepresented in the infected hACE2 mice in Metascape and Reactome pathway analysis (**Figure 3A, and Supplementary Figure 5C**). Interestingly, in addition to immune signaling, both analyses showed the upregulation of a large number of genes related to cell cycle and cell cycle checkpoints pathways in infected lungs (**Figure 3A and Supplementary Figure 5C**). The TRRUST analysis (transcriptional regulatory network analysis, http://www.grnpedia.org/trrust) indicate that *Irf1*, *Stat1*, and *Nfkb1/Rela* are the topmost predicted transcription factors regulating the genes upregulated in hACE2 mice lungs (**Figure 3A, right panel**). A vast number of immunity-related genes including chemokines/cytokines (*Cxcl9, Cxcl10, Cxcl11, Ccl1, Ccl7*), interleukins (*IL6, IL1b, IL12, IL24*), Toll-like receptors, C-type lectins receptors, tumor necrosis factor (*Tnf*) associated genes, *Ifng*, interferon-stimulated genes (ISGs) (*Oas* family genes, *Ifi* genes *Gbp* family genes, *Irf* genes) were upregulated in SARS-CoV-2 infected hACE2 mice lungs (**Supplementary Figure 5D-5F**). The genes and surface receptors related to T-Cells, NK cells, macrophages, and neutrophils were induced in the lungs of hACE2 mice infected with SARS-CoV-2 (**Supplementary Figure 5G and 5H**) suggesting a massive infiltration of immune cells into the lungs confirming the immunohistochemistry (IHC) data shown in Figure 2. Several genes related to neutrophil degranulation were also upregulated in the lungs of hACE2 mice infected with SARS-CoV-2 (**Supplementary Figure 5I**). Thus. a massive cytokine storm is induced in the lungs of SARS-CoV-2-infected mice, which would have contributed to extensive pathology observed in these lungs.

**Figure 3.**
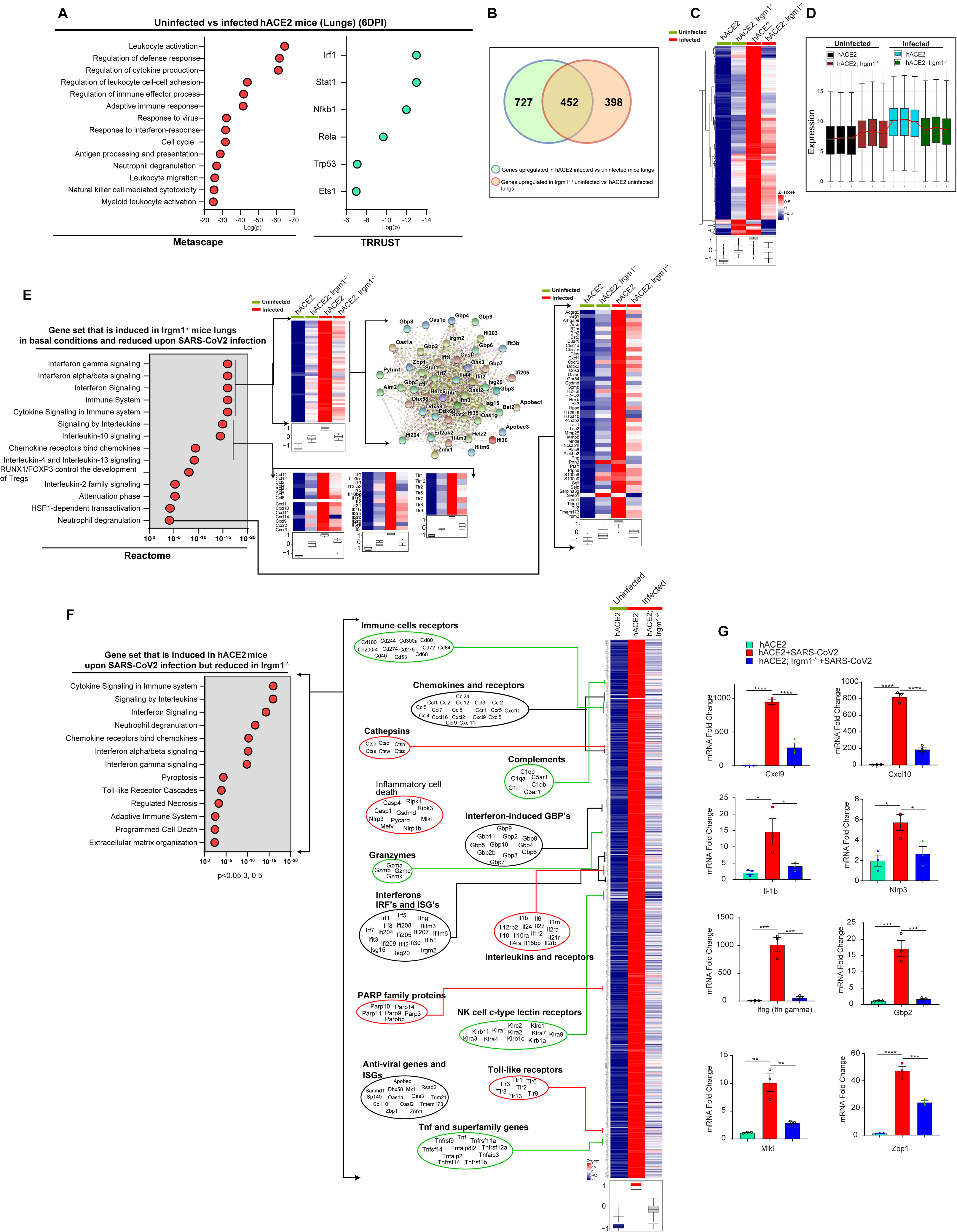
SARS-CoV-2 infection-driven cytokine storm is blunted in hACE2 mice lacking Irgm1. **A.** Gene Ontology-based pathway analysis (using Metascape) performed with geneset that is upregulated in SARS-CoV-2 infected hACE2 mice compared to uninfected hACE2 mice at 6 dpi in lungs (p<0.05, 3 folds, n=3). Graph depicts top enriched pathways. TRRUST analysis from the transcriptome induced in SARS-CoV-2 infected hACE2 mice compared to uninfected hACE2 mice at 6 dpi (p<0.05, 3 folds, n=3) to predict transcription factors that drive the gene expression. **B.** Venn diagram shows the overlap between genes that are upregulated upon SARS-CoV-2 infection in hACE2 mice lungs (p < 0.05; 3 folds; n=3) vs induced in *Irgm1* knockout lungs in basal conditions (p < 0.05; 1.5 folds; n=3) at 6 dpi. **C.** Hierarchical clustering of a set of differentially expressed genes (p<0.05, Wald Chi-Squared test, n=3, 6 dpi lungs) that are upregulated (1.4 folds) in hACE2; *Irgm1^−/−^* mice (compared to uninfected hACE2 mice) in basal conditions and at the same time are suppressed (>25%) in the SARS-CoV-2 infected hACE2; *Irgm1^−/−^* mice (compared to uninfected hACE2 mice). **D.** Box whisker distribution plot with normalized log expression of the geneset that is significantly upregulated in lungs of uninfected hACE2; *Irgm1^−/−^* mice (compared to uninfected hACE2; 1.4 fold, p<0.05, Wald Chi-Squared test, n=3) and at the same time suppressed in lungs of SARS-CoV-2 infected hACE2; *Irgm1^−/−^* mice (compared to infected hACE2; 0.75 fold, p<0.05, Wald Chi-Squared test, n=3) at 6 dpi. **E.** Pathway analysis (Reactome) using geneset (p<0.05, Wald Chi-Squared test, n=3, 6 dpi) that is upregulated (1.4 folds) in hACE2; *Irgm1^−/−^* mice (compared to uninfected hACE2 mice) in basal conditions and at the same time are suppressed (>25%) in the SARS-CoV-2 infected hACE2; *Irgm1^−/−^* mice (compared to uninfected hACE2 mice). Genes related to top pathways (interferon signaling, immune system, cytokine response, and neutrophil degranulation) are shown using heatmaps and/or STRING networks. **F.** Pathway analysis (Reactome) using geneset (p<0.05, Wald Chi-Squared test, n=3, 6 dpi) that is upregulated (3 folds) in the SARS-CoV-2 infected hACE2 mice (compared to uninfected hACE2 mice) and at the same time are suppressed (>50%) in the SARS-CoV-2 infected hACE2; *Irgm1^−/−^* mice. The genes related to top pathways are depicted. **G.** qRT-PCR validation of RNA-Seq data with total RNA isolated from lungs of uninfected hACE2 and SARS-CoV-2 infected hACE2, hACE2; *Irgm1^−/−^* mice at 6 dpi. Mean ± SEM, n=3, *p < 0.05, **p < 0.005, ***p < 0.0005 and ****p < 0.00005, One-way ANOVA (Tukey’s multiple comparison test).

IRGM/Irgm1 is a master suppressor of type 1 interferon response (Jena et al, 2020). First, we examined whether hACE2; *Irgm1^−/−^* mice like parental *Irgm1^−/−^* mice (Jena et al, 2020) show sustained pro-inflammatory and anti-viral responses. An analysis of significantly upregulated genes in hACE2; *Irgm1^−/−^* mice compared to hACE2 mice revealed the presence of pathways associated with immune responses, including type 1 interferon response, interleukin response, and complement pathway (**Supplementary Figure 6A-6C; Supplementary data set 2**). Thus, hACE2; *Irgm1^−/−^* mice are similar to the parental *Irgm1^−/−^* strains in terms of sustained preexisting IFN-I response.

Interestingly, about 452 genes (>50% genes) genes that are induced (1.5 folds, p<0.05) in SARS-CoV-2 infected hACE2 mice lungs (compared to uninfected hACE2 mice) were common to those that are induced (1.5 folds, p<0.05) in hACE2; *Irgm1^−/−^* mice lungs in basal conditions (compared to uninfected hACE2 mice) (**Figure 3B**). Further, a large number of genes that are related to pathways such as cytokine signaling (90 genes), interferon signaling (45 genes), neutrophil degranulation (36) and adaptive immune response (76 genes) were overlapping between these two transcriptomes (**Supplementary Figure 6D**). Consistently, the predicted top transcription factors (*Irf1*, *Stat1*, and *Nfkb1/Rela*) for these transcriptomes were also found to be the same (**Supplementary Figure 6E and Figure 3A**). This data show that hACE2; *Irgm1^−/−^* mice already possess the host antiviral response that is induced by SARS-CoV-2, which may explain their strong antiviral phenotype noted above.

The GO-based pathway analysis was performed on a set of genes that were upregulated under basal conditions in hACE2; *Irgm1^−/−^* mice (compared to uninfected hACE2 mice), and at the same time were suppressed in hACE2; *Irgm1^−/−^* mice upon infection with SARS-CoV-2 (compared to uninfected hACE2 mice) (**Figure 3C and 3D, Supplementary Dataset 3**). The data show that all the inflammatory/antiviral pathways that were induced in basal conditions in hACE2; *Irgm1^−/−^* mice were further upregulated in SARS-CoV-2 infected hACE2 mice lungs (**Figure 3E**). Whereas these responses were attenuated in hACE2; *Irgm1^−/−^* mice during SARS-CoV-2 infection (**Figure 3E**).

We also performed pathway analysis with the geneset that is induced in hACE2 mice lungs upon SARS-CoV-2 infection (compared to uninfected hACE2 mice lungs) and at the same time suppressed in lungs of SARS-CoV-2 infected hACE2; *Irgm1^−/−^* mice lungs (**Figure 3F, Supplementary Dataset 4**). We found that a substantial number of genes belonging to pathways of interferon responses (*Ifng, Irf1, Irf5, Irf7, Irf8, Isg15, Ifitm3, Ifi30, Ifi204, Ifi208, Ifih1, Apobec1, Gbp2, Gpb3, etc) cytokine/chemokine* response (*Il1b, Il6, Il24, Il27, Ccl1, Ccl2, Ccl3, Ccl24, Cxcl2, Cxcl9,* etc) neutrophil degranulation, immune cells receptors, pyroptosis (*Nlrp3, Pycard, Casp1, Gsdmd*), T-cell/NK cell/macrophage receptors, TNF family genes *(Tnf, Tnsf9, Tnsf14, Tnsf11a),* genes involved in necrosis, programmed cell death, and tissue damage that were considerably upregulated (3 folds, p<0.05) in hACE2 mice lungs were significantly suppressed (0.5 folds, p<0.05) in hACE2; *Irgm1^−/−^* mice (**Figure 3F, Supplementary Dataset 4**). The RNA-seq data is validated using qRT-PCR with some of the sentinel pro-inflammatory genes (**Figure 3G**). Taken together, the data show that SARS-CoV-2 mediated cytokine storm is attenuated in *Irgm1*-depleted mice.

### Enhanced SARS-CoV-2 infection, immunopathology and cytokine storm in lungs of hACE2; Ifnar1^−/−^ mice

To further validate the protective function of type 1 IFN response against SARS-CoV-2 infection, we developed hACE2 expressing *Ifnar1* knockout mice by crossing hACE2 and *Ifnar1^−/−^* mice (**hereafter, hACE2; *Ifnar1^−/−^***; hemizygous for hACE2 gene and homozygous knockout for *Ifnar1*). The hACE2 and hACE2; *Ifnar1^−/−^* mice were infected with low dose (5X10^4^) SARS-CoV-2 (clade 19A strain) and the symptoms were monitored. The hACE2; *Ifnar1^−/−^* mice lost weight significantly faster than hACE2 mice (**Figure 4A**). Rapid hypothermia (**Figure 4B**) and significantly faster appearance of severe clinical symptoms were observed in hACE2; *Ifnar1^−/−^* mice (**Figure 4C; Supplementary Video 3**). Consistently, hACE2; *Ifnar1^−/−^* mice reached the euthanasia conditions significantly faster than hACE2 mice (**Figure 4D**).

**Figure 4.**
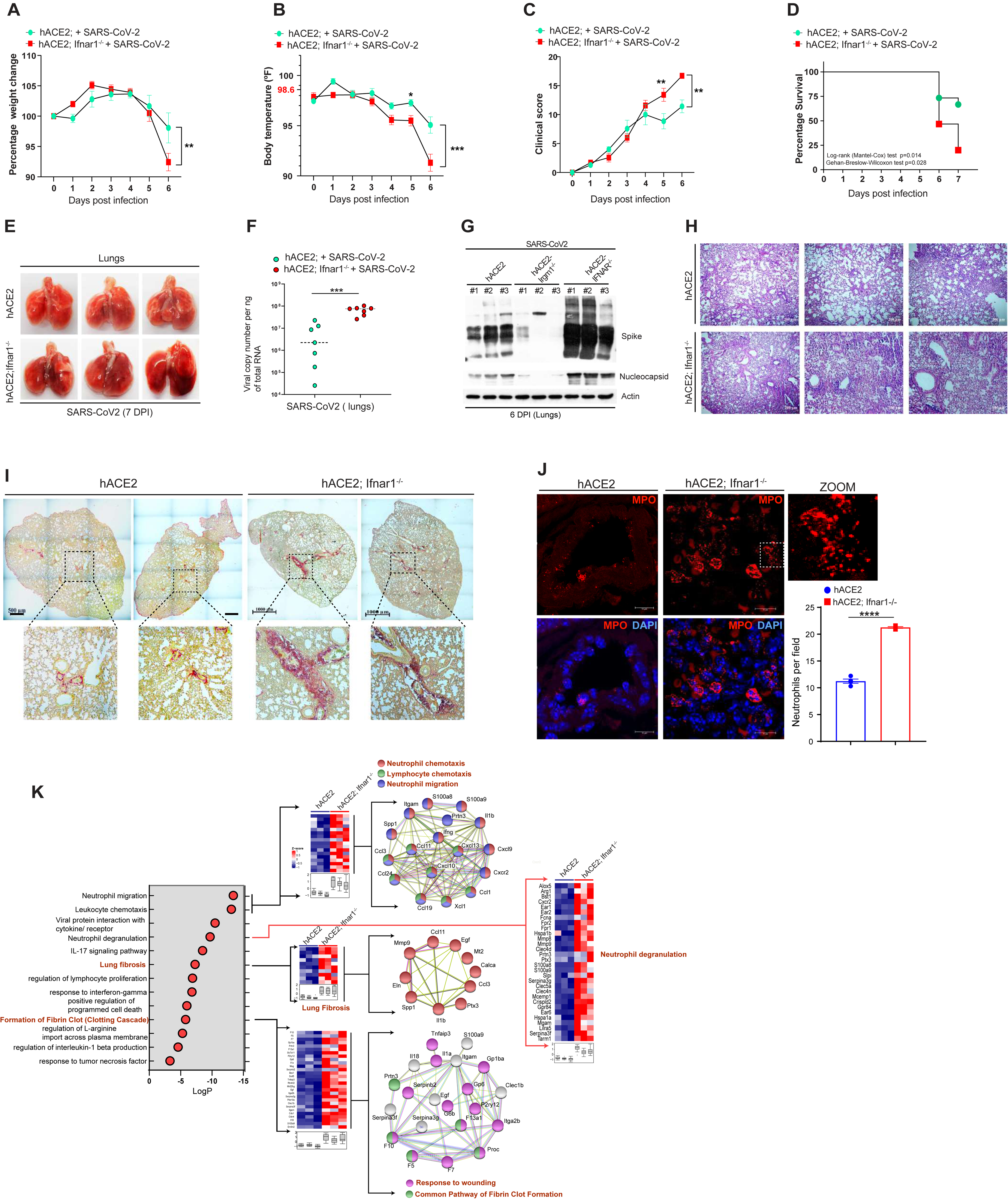
Enhanced SARS-CoV-2 infection, immunopathology, and cytokine storm in lungs of hACE2; Ifnar1^−/−^ mice. **A-C.** Graphs depict **(A)** percentage change in body weight, **(B)** core body temperature (in °F) and **(C)** clinical scores of SARS-CoV-2 infected hACE2 (n=7) and hACE2; *Ifnar1*^−/−^ (n=9) mice for 6 days (mean ± SEM, *p < 0.05, **p < 0.01 and ***p < 0.001, Two-way ANOVA). **D.** Kaplan–Meier survival graph showing percentage survival of hACE2 and hACE2; *Ifnar1*^−/−^ mice during the course of SARS-CoV-2 infection [*p = 0.014, Log-rank (Mantel-Cox) test, *p = 0.028, Gehan-Breslow-Wilcoxon test]. **E.** Lung images depicting gross pathology in hACE2 and hACE2; *Ifnar1*^−/−^ mice infected with SARS-CoV-2 (5X10^4^ PFU/mice) at 7 dpi. **F.** Assessment of viral copy number in total RNA isolated from lungs of SARS-CoV-2 infected hACE2 and hACE2; *Ifnar1*^−/−^ mice at 6, 7 dpi (n=7). Mean ± SEM, ***p < 0.001, Student’s unpaired t-test. **G.** Western blot analysis of Spike and NC protein expression in lung lysates of SARS-CoV-2 infected hACE2, hACE2; Irgm1^−/−^ and hACE2; *Ifnar1*^−/−^ mice at 6 dpi. **H.** Representative hematoxylin & eosin stained lung section images of SARS-CoV-2 infected hACE2 and hACE2; *Ifnar1*^−/−^ mice. **I.** Representative picrosirius red stained lung images of SARS-CoV-2 infected hACE2 and hACE2; *Ifnar1*^−/−^ mice showing collagen deposition (in red). **J.** Representative confocal images of MPO (activated neutrophils) immunostained lung sections of SARS-CoV-2 infected hACE2 and hACE2; *Ifnar1*^−/−^ mice. Bar graphs depict average number of MPO positive cells per field (10 fields from each experiment, n=4 experiments, mean ± SEM, ****p < 0.0001, Student’s unpaired t-test). **K.** Gene Ontology-based pathway analysis (using Reactome) performed with geneset that is upregulated in lungs of SARS-CoV-2 infected hACE2; *Ifnar1*^−/−^ mice (compared to infected hACE2; 2 folds, p<0.05, n=3) at 7 dpi. Graph depicts top enriched pathways. Heatmaps and STRING functional protein-protein association networks are used to depict the genes and pathways involved in neutrophil migration, leukocyte chemotaxis, neutrophil degranulation, lung fibrosis, and formation of fibrin clot.

Grossly, lungs of hACE2; *Ifnar1^−/−^* mice appeared more inflamed than hACE2 mice (**Figure 4E**). Consistently, the SARS-CoV-2 mRNA and protein (spike and NC) levels were significantly higher in the lungs of hACE2; *Ifnar1^−/−^* mice than in hACE2 mice (**Figure 4F and 4G**). The histopathology results showed a higher degree of alveolar space consolidation, alveolar septa thickening, parenchymal collapse, and increased infiltration of immune cells in hACE2; *Ifnar1^−/−^* mice (**Figure 4H, Supplementary Figure 7A**). The PSR staining of the lung section depicted fibrosis in hACE2; *Ifnar1^−/−^* mice even at 6 DPI (**Figure 4I**). Consistently, hACE2; *Ifnar1^−/−^* mice had significantly higher neutrophil infiltration into their lungs than hACE2 mice (**Figure 4J**), however, macrophage infiltration was not different (Data not shown).

RNA-seq analysis was performed to determine the genes and pathways responsible for the dramatically increased lung pathology in hACE2; *Ifnar1^−/−^* mice compared with hACE2 mice. PCA plot suggests a distinct transcriptional signature of all the groups (**Supplementary Figure 7B**). As expected, pathway analysis of genes that are suppressed (0.5 folds, p<0.05) in lungs of hACE2; *Ifnar1^−/−^* mice (compared to hACE2 mice) showed the type 1 IFN response as one of the top pathways significantly suppressed (**Supplementary Figure 7C**).

We then performed pathway analysis with a set of genes (**Supplementary Dataset 5**) significantly upregulated (>2 folds, p<0.05) in hACE2; *Ifnar1^−/−^* mice lungs as compared to the hACE2 mice. The pathways related to immune cell migration, neutrophil degranulation, IL-17 signaling pathway, IL-1β signaling pathways, Interferon-gamma signaling, and TNFα response were overrepresented (**Figure 4K**) suggesting a massive pro-inflammatory response in hACE2; *Ifnar1^−/−^* mice lungs. The top 50 induced genes induced in hACE2; *Ifnar1^−/−^* mice were found to have a function in immune cell migration, cytokine/chemokine response, neutrophil degranulation, macrophage antimicrobial response, coagulation factors, extracellular matrix remodeling, etc. (**Supplementary Figure 7D and Supplementary Dataset 6**). Further, the genes and GO terms related to positive regulation of programmed cell death, lung fibrosis, and formation of fibrin clots were significantly upregulated in the transcriptome of hACE2; *Ifnar1^−/−^* mice (**Figure 4K**). Taken together, the RNA-seq analysis provided a molecular basis for amplified pathology in the lungs of hACE2; *Ifnar1^−/−^* mice compared to hACE2 mice.

### SARS-CoV-2 infection, cytokine storm, and immune cell infiltration in hACE2; Ifnar1^−/−^ mice brain

Next, we asked whether the higher magnitude of the infection in hACE2; *Ifnar1^−/−^* mice increased their vulnerability to SARS-CoV2 neuroinvasion in the brain. We found that where none of the seven hACE2 mice in this experiment showed significant viral mRNA levels, an excessive amount of SARS-CoV-2 mRNA was detected in the brains of hACE2; *Ifnar1^−/−^* mice. (**Figure 5A**). Furthermore, Western blot analysis shows a significant level of NC in brain lysates of hACE2; *Ifnar1^−/−^* mice, while none is detected in hACE2 mice. (**Figure 5B**). To ascertain, we infected a total of eleven hACE2; *Ifnar1^−/−^* mice with SARS-CoV-2 and performed Western blotting with NC. The nucleocapsid protein was detected in all (100%) of the mice with ten mice having very high NC levels (**Supplementary Figure 7E**). This result is in striking contrast with the rate of infection (23.5%, 4/17) in hACE2 mice brains tested in the previous experiment (**Supplementary Figure 3G**). The expression of ACE2 was found to be almost the same in hACE2 and hACE2; *Ifnar1^−/−^* mice (**Supplementary Figure 7F**). In immunofluorescence assays, substantially high SARS-CoV-2 infection was observed in hACE2; *Ifnar1^−/−^* mice (**Figure 5C and 5D**). The NC was detected in vesicles-like structure in cells and was found to be co-localized with EEA1, an early endosome marker, suggesting a cellular invasion and replication of SARS-CoV-2 in the brain of hACE2; *Ifnar1^−/−^* mice (**Supplementary Figure 7G**).

**Figure 5.**
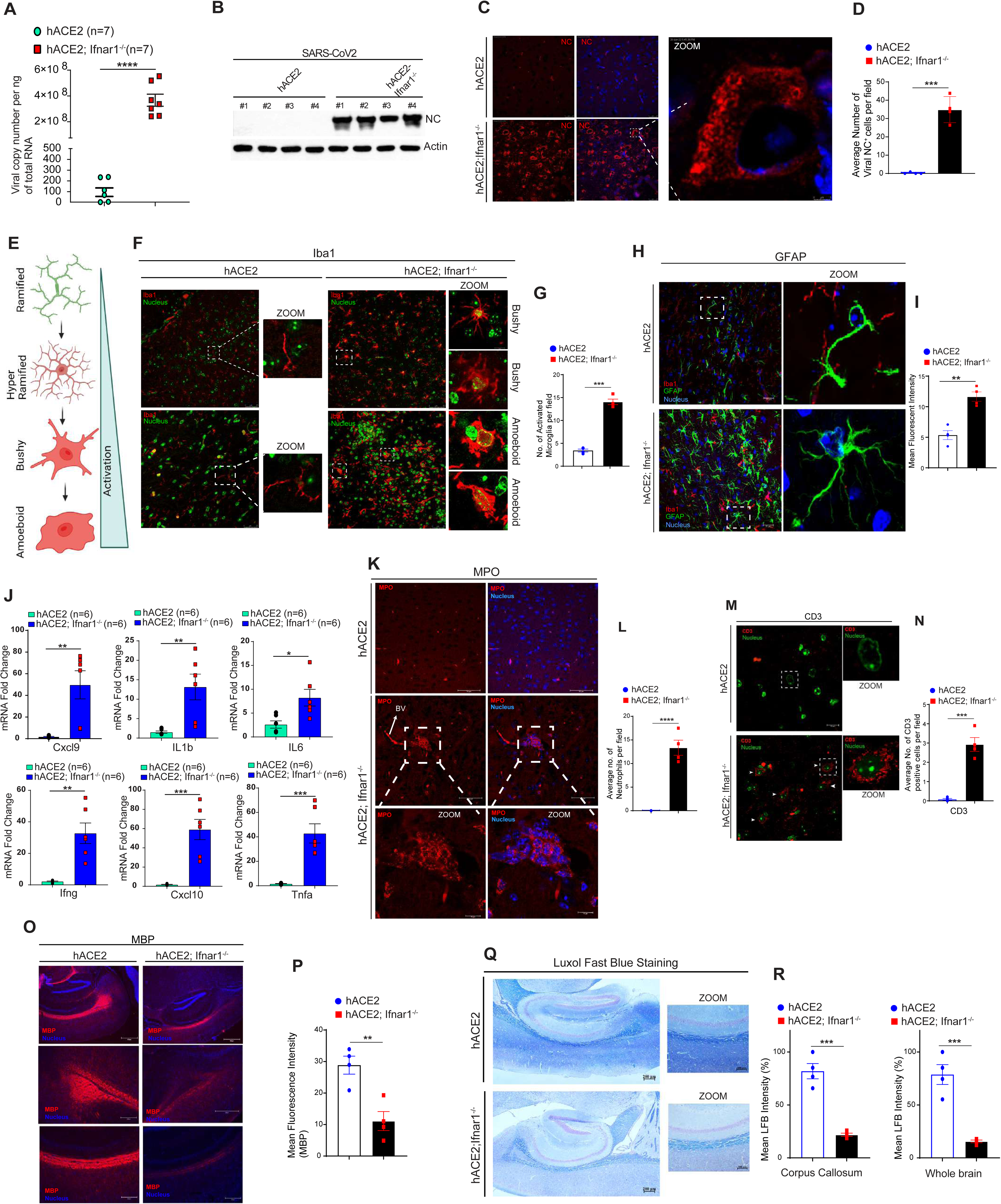
Enhanced SARS-CoV-2 infection, cytokine storm, and immune cell invasion in hACE2; Ifnar1^−/−^ mice brain. **A.** Viral copy number analysis in total RNA isolated from the brain of SARS-CoV-2 infected hACE2 and hACE2; *Ifnar1*^−/−^ mice at 6, 7 dpi (n=7). Mean ± SEM, ****p < 0.0001, Student’s unpaired t-test. **B.** Western blot analysis of NC protein expression in brain lysates of SARS-CoV-2 infected hACE2 and hACE2; *Ifnar1*^−/−^ mice at 6 dpi. **C-D. (C)** Representative confocal images of NC immunostained brain sections of SARS-CoV-2 infected hACE2 and hACE2; *Ifnar1*^−/−^ mice at 7 dpi. Nucleus stained with DAPI. Right panel shows zoomed image. (**D)** Bar graphs shows average number of NC positive cells per field (10 fields from each experiment, n=4 experiments, mean ± SEM, ***p < 0.001, Student’s unpaired t-test). **E.** Schematic showing microglial remodeling, its different activation states from ramified (resting, quiescent) to amoeboid (fully activated, phagocytic) state. Image created using Biorender.com. **F-G.** (**F**) Representative confocal images of IBA1 immunostained brain sections of SARS-CoV-2 infected hACE2 and hACE2; *Ifnar1*^−/−^ mice (7 dpi). Nucleus stained with DAPI. Right panels show zoomed images of microglial cells marked in the inset showing their different activation state. (**G**) Bar graphs showing average number of IBA1 positive cells per field (10 fields from each experiment, n=4 experiments, mean ± SEM, ***p < 0.001, Student’s unpaired t-test). **H-I.** (**H**) Representative confocal images of GFAP (Astrocytes) immunostained brain sections of SARS-CoV-2 infected hACE2 and hACE2; *Ifnar1*^−/−^ mice (7 dpi). Nucleus stained with DAPI. Right panels showing zoomed image. (**I**) Bar graphs showing mean fluorescence intensity of GFAP in brain (10 fields from each experiment, n=4 experiments, mean ± SEM, ***p < 0.001, Student’s unpaired t-test). **J.** qRT-PCR analysis for indicated cytokines genes with total RNA isolated from brain of SARS-CoV-2 infected hACE2 and hACE2; *Ifnar1*^−/−^ mice (6 dpi). (n=6, Mean ± SEM *p < 0.05, **p < 0.01, ***p < 0.001, Student’s unpaired t-test). **K-L.** (**K**) Representative confocal images of MPO (activated neutrophils) immunostained brain sections of SARS-CoV-2 infected hACE2 and hACE2; *Ifnar1*^−/−^ mice (7 dpi). Bottom panels showing zoomed image. (**L**) Bar graphs showing average number of MPO positive cells per field (10 fields from each experiment, n=4 experiments, mean ± SEM, ****p < 0.0001, Student’s unpaired t-test). **M-N.** (**M**) Representative confocal images of CD3 (T cells) immunostained brain sections of SARS-CoV-2 infected hACE2 and hACE2; *Ifnar1*^−/−^ mice (7 dpi). Right panels showing zoomed image. (**N**) Bar graphs showing average number of CD3 positive cells per field (10 fields from each experiment, n=4 experiments, mean ± SEM, ***p < 0.001, Student’s unpaired t-test). **O-P.** (**O**) Representative images of MBP (Myelin Basic Protein) immunostained brain (corpus callosum) sections of SARS-CoV-2 infected hACE2 and hACE2; *Ifnar1*^−/−^ mice (7 dpi). (**P**) Bar graphs showing mean fluorescence intensity of MBP in brain (10 fields from each experiment, n=4 experiments, mean ± SEM, **p < 0.001, Student’s unpaired t-test). **Q-R.** (**Q**) Representative Luxol Fast Blue (LFB) stained brain images of SARS-CoV-2 infected hACE2 and hACE2; *Ifnar1*^−/−^ mice (7 dpi). (**R**) Bar graph depicts the mean Luxol fast blue (LFB) intensity (in percentage) in corpus callosum and whole brain. (10 fields from each experiments, n=4 experiment, mean ± SEM, ***p < 0.001, Student’s unpaired t-test).

A universal component of neuroinflammation is the hyperactivation of microglial and astrocytes, which is implicated in the progression of neurological disorders (Glass et al, 2010). The reactive and inflammatory microglial cells have bushy and amoeboid morphology compared to the ramified shape of the resting microglial (**Figure 5E**). Microgliosis (Reactive microglial cells) was observed throughout the hACE2; Ifnar1^−/−^ mice brain especially in the olfactory lobe, pre-frontal cortex, cortex, hippocampus, thalamus, and medulla region (**Figure 5F and 5G**), indicating a hyperinflammatory microenvironment. Whereas normal ramified microglial cells were observed throughout the hACE2 mice brains (**Figure 5F**). Reactive astrocytes are characterized by high-level expression of intermediate filament glial fibrillary acidic protein (GFAP, a marker for astrocytes) and an increase in the number and length of GFAP-positive processes(Escartin et al, 2021). A significantly higher number of reactive astrocytes (Astrogliosis) were observed in the corpus callosum, olfactory lobe, and prefrontal cortex region of hACE2; *Ifnar1^−/−^* mice brains (**Figure 5H and 5I**). Consistently, dramatically enhanced levels of potent pro-inflammatory molecules including chemokines (such as *Cxcl9, Cxcl10)*, *Ifng*, *Tnfa*, and interleukins (*Il6 and Il1b*) were observed in hACE2; *Ifnar1^−/−^* mice brains (**Figure 5J**).

Since the cytokine response was so high in hACE2; *Ifnar1^−/−^* mice brain, we suspected infiltration of circulatory immune cells. Hence, we measured the levels of neutrophils and T-cells infiltrating the brain. MPO puncta positive neutrophils (activated neutrophils) were present throughout the hACE2; *Ifnar1^−/−^* mice brain, especially near the blood vessels (**Figure 5K and 5L**), as opposed to no infiltration in hACE2 mice brains (**Figure 5K and 5L**). A significant number of CD3-positive T-cells was also observed in the brains of hACE2; *Ifnar1^−/−^* mice but none was detected in hACE2 mice (**Figure 5M and 5N**).

Hyperinflammation is the most common cause of demyelination of neurons in the brain, commonly occurring during multiple sclerosis disorder or viral infections (Teeling & Perry, 2009). While SARS-CoV-2 is typically transmitted through the respiratory tract, there are evidence that the virus invades the central nervous system. Case reports describe SARS-CoV-2 patients with demyelinating lesions in the optic nerve, spinal cord, brain, and, suggesting possible implications for neuroimmune disorders such as multiple sclerosis and others (MacDougall et al, 2022; Moore et al, 2021; Palao et al, 2020; Sarwar et al, 2021; Zanin et al, 2020). Thus, we assessed the status of myelination in SARS-CoV-2-infected mice brains using conventional markers, myelin basic protein (MBP), and luxol fast blue (LFB) staining. There was dramatic demyelination of neurons in the brains (especially, the corpus callosum region) of SARS-CoV-2 infected hACE2; *Ifnar1^−/−^* mice as compared to hACE2 mice (**Figure 5O-5R, Supplementary Figure 7H**).

Taken together, the data show that deficiency of type 1 IFN response enhances susceptibility to SARS-CoV-2 neuroinvasion and subsequent neuroinflammation (characterized by microgliosis and astrogliosis) and immunopathology in the brain.

### Irgm1 depletion is more protective than IFN-α treatment against SARS-CoV-2 infection

Type 1 IFNs administered before a viral infection or post-infection but before the inflammatory response peaks have been shown to offer a high level of protection (Bessiere et al, 2021; Channappanavar et al, 2019; Lee et al, 2021; Sabit et al, 2018; Sodeifian et al, 2022). Several studies even reported the beneficial effect of IFN-α in the treatment of SARS-CoV-2-infected patients (Bhushan et al, 2021; Pandit et al, 2021; Pereda et al, 2020a; Pereda et al, 2020d; Rahmani et al, 2020; Zhou et al, 2020a; Zhou et al, 2020b).

We next asked how *Irgm1* depletion compares to IFN-α therapy in protecting against SARS-CoV-2 infection. The hACE2 mice were administrated with IFN-α for three consecutive days before SARS-CoV-2 infection as depicted in the schematic (**Figure 6A**). Both the hACE2; *Irgm1^−/−^* mice and IFN-α treated hACE2 mice showed significantly milder clinical symptoms than control hACE2 mice (**Figure 6B**). Consistent with our previous results, the gross lung pathology of hACE2; *Irgm1^−/−^* mice were milder compared to hACE2 mice (**Figure 6C**). However, IFN-α treated hACE2 mice showed gross lung pathology (**Figure 6C**).

**Figure 6.**
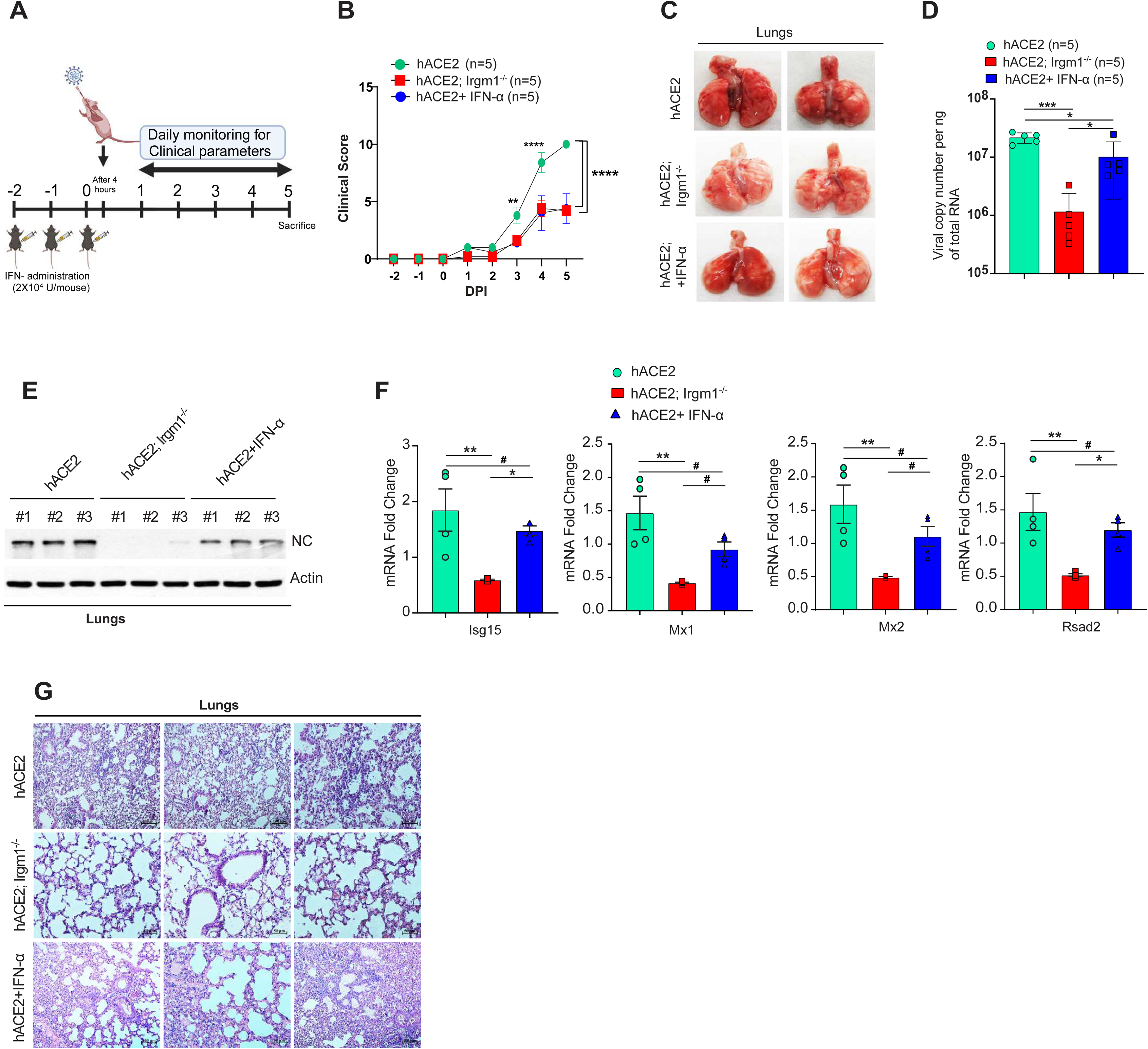
Irgm1 depletion is more protective to SARS-CoV-2 infection than IFN-α treatment. **A.** Schematic representation of the experimental design depicting IFN-α treatment (20,000U/mouse) starting 2 days before the SARS-CoV-2 (5X10^4^ PFU) infection. Image created using Biorender.com. **B.** Graph showing total clinical scores of hACE2, hACE2; *Irgm1^−/−^* and hACE2 + IFN-α treated mice infected with SARS-CoV-2 for 5 days. n=5, Mean ± SEM, **p < 0.01, ****p < 0.0001, (Two-way ANOVA, Tukey’s multiple comparison test). **C.** Lung images showing gross pathology in hACE2, hACE2; *Irgm1^−/−^* and hACE2 + IFN-α treated mice infected with SARS-CoV-2 at 5 dpi. **D.** Assessment of viral copy number in total RNA isolated from lungs of SARS-CoV-2 infected hACE2, hACE2; *Irgm1^−/−^* and hACE2 + IFN-α treated mice at 5 dpi (n=5). Mean ± SEM, *p < 0.05, ***p < 0.001, One-way ANOVA (Tukey’s multiple comparison test). **E.** Western blot analysis of NC protein expression in lung lysates of SARS-CoV-2 infected hACE2, hACE2; *Irgm1^−/−^* and hACE2 + IFN-α treated mice at 5 dpi. **F.** qRT-PCR analysis for indicated ISGs with total RNA isolated from lungs of SARS-CoV-2 infected hACE2, hACE2; Irgm1^−/−^ and hACE2 + IFN-α treated mice (5 dpi). n=4, mean ± SEM *p < 0.05, **p < 0.01, ^#^ ns, One-way ANOVA (Tukey’s multiple comparison test). **G.** Representative hematoxylin & eosin stained lung images of SARS-CoV-2 infected hACE2, hACE2; *Irgm1^−/−^* and hACE2 + IFN-α treated mice at 5 dpi.

A mild but significant reduction in viral load was observed in IFN-α treated group compared to control hACE2 mice (**Figure 6D**). However, the SARS-CoV-2 mRNA levels in IFN-α treated group was found to be significantly higher than hACE2; *Irgm1^−/−^* mice (**Figure 6D**). Similar results were obtained in the immunoblotting experiment using SARS-CoV-2 NC antibody (**Figure 6E**) suggesting that the effect of Irgm1 depletion is much stronger than IFN-α treatment. In agreement with a low viral infection in hACE2; *Irgm1^−/−^* mice, the expression of ISGs was significantly less compared to the control. In contrast, the IFN-α treatment group showed consistently enhanced expression of the ISGs (**Figure 6F**). Histology data showed reduced pathology including alveolar consolidation and septal thickening in hACE2; *Irgm1^−/−^* mice compared to both control hACE2 mice and the IFN-α treated hACE2 mice (**Figure 6G**).

The results show that *Irgm1* depletion could provide better protection than IFN-α treatment against SARS-CoV-2 infection without provoking severe cytokine responses.

### Inhibition of STING and RIPK2 in hACE2; Irgm1^−/−^ mice enhances their susceptibility to SARS-CoV-2 infection

We showed that upon *Irgm1* depletion, the cGAS-STING-IRFs and NODs-RIPK2-NFκB signaling are activated to induce type 1 IFN response (Jena et al, 2020; Mehto et al, 2022). We next examined whether type 1 IFN response is responsible for protective phenotypes observed in hACE2; *Irgm1^−/−^* mice by inhibiting the STING and RIPK2 (**Supplementary Figure 8A and 8G**).

The inhibition of STING using H-151 (Gong et al, 2021; Haag et al, 2018; Pan et al, 2021) in hACE2; *Irgm1^−/−^* mice enhanced the clinical symptoms associated with SARS-CoV-2 infection (**Supplementary Figure 8B**). Further, the H-151 treated mice lungs showed significantly higher viral mRNA levels and nucleocapsid protein levels than hACE2; *Irgm1^−/−^* mice (**Supplementary Figure 8C and 8D**). Consistent with higher viral loads in the lungs of H-151, the levels of sentinels ISG’s (*Mx1 and Isg15*) were found to be significantly upregulated in these mice compared to hACE2; *Irgm1^−/−^* mice (**Supplementary Figure 8E and 8F**). GSK583 is a potent inhibitor of RIPK2 and can suppress NF-κB response. Recently, we found that GSK583 could suppress NF-κB-dependent interferon and cytokine response in IRGM-depleted human cells and *Irgm1^−/−^* mice (Mehto et al, 2022). GSK583 treated hACE2; *Irgm1^−/−^* mice (**Supplementary Figure 8G**) showed clinical symptoms similar to hACE2 mice indicating suppression of protective effects (**Supplementary Figure 8H**). In agreement, significantly higher levels of viral mRNA (**Supplementary Figure 8I**), nucleocapsid proteins (**Supplementary Figure 8J**), and ISG’s (*Mx1 and Isg15*) (**Supplementary Figure 8K and 8L**) were observed in GSK583 treated hACE2; *Irgm1^−/−^* mice than control hACE2; *Irgm1^−/−^* mice. The H&E staining of lung sections shows higher pathological changes including alveolar consolidation and septal thickening in GSK583 treated mice lungs compared to hACE2; *Irgm1^−/−^* mice (**Supplementary Figure 8M**).

Altogether, the results show that heightened STING and RIPK2-dependent immunity in hACE2; *Irgm1^−/−^* mice protect them from severe SARS-CoV-2 infection.

### Depletion of anti-viral factors in Irgm1^−/−^ BMDMs increases their susceptibility to SARS-CoV-2 infection

Irgm1 depletion induces several well-known ISGs and potent antiviral factors such as *Mx1, Isg15, Oas1, Ifitm3,* and *Rsad2* (Viperin) (Jena et al, 2020; Nath et al, 2021). We knock down these anti-viral factors in bone marrow-derived macrophages (BMDMs) isolated from hACE2; *Irgm1^−/−^* mice and infected with SARS-CoV-2 (MOI, 0.5) for 16h. The hACE2; *Irgm1^−/−^* BMDMs were found to be resistant to SARS-CoV-2 infection (**Supplementary Figure 9A**). The depletion of *Mx1, Isg15, Oas1, Ifitm3, and Rsad2* (Viperin) in hACE2; *Irgm1^−/−^* BMDMs (**Supplementary Figure 9B)** significantly enhanced the susceptibility of SARS-CoV-2 infection (**Supplementary Figure 9A)**. The data suggest that the heightened expression of several ISGs and anti-viral host factors protects *Irgm1*-depleted mice from SARS-CoV-2 infection.

### Irgm1^−/−^ Ifnar1^−/−^ double knockout mice were not protected from SARS-CoV-2 infection and displayed enhanced immunopathology

Next, we generated hACE2 expressing *Irgm1* and *Ifnar1* double knockout mice (**hereafter, hACE2; *Irgm1^−/−^ Ifnar1^−/−^***, hemizygous for hACE2 gene and homozygous knockout for *Irgm1 and Ifnar1*). The three groups of mice, hACE2, hACE2; *Irgm1^−/−^*, and hACE2; *Irgm1^−/−^ Ifnar1^−/−^* were infected with 5X10^4^ PFU of SARS-CoV-2 and symptoms were monitored for 6 Days before the mice were sacrificed (**Figure 7A**). Where hACE2; *Irgm1^−/−^* mice showed minor signs of infection, the hACE2; *Irgm1^−/−^ Ifnar1^−/−^* displayed clinical symptoms similar or more exacerbated than hACE2 mice on 6 DPI (**Figure 7B**). In contrast to hACE2; *Irgm1^−/−^* mice, rapid hypothermia and loss of weight were observed in hACE2 and hACE2; *Irgm1^−/−^ Ifnar1^−/−^* mice (**Figure 7C-7D**). The lungs of hACE2; *Irgm1^−/−^ Ifnar1^−/−^* mice were highly inflamed with visible lesions, which was in striking contrast to the seemingly normal lungs of hACE2; *Irgm1^−/−^* mice (**Figure 7E**). Consistently, SARS-CoV-2 mRNA and protein levels were substantially elevated in hACE2; *Irgm1^−/−^ Ifnar1^−/−^* as compared to hACE2; *Irgm1^−/−^* mice (**Figure 7F and 7G**). Acute lung injury leads to increased damage of pulmonary microvasculature and alveolar epithelium resulting in enhanced lung permeability and pulmonary edema. The lungs’ vascular and epithelial permeability can be measured using permeability to Evans blue dye. An increased Evans blue staining was observed in the lungs of hACE2 and hACE2; *Irgm1^−/−^ Ifnar1^−/−^* mice suggesting a higher level of lung damage, vascular/epithelial permeability, and pulmonary edema (**Figure 7H**). In histopathology analysis, similar to hACE2 mice, hACE2; *Irgm1^−/−^ Ifnar1^−/−^* mice lungs showed considerable lung damage with significant consolidation, alveolar septa thickening, parenchymal collapse, and infiltration of immune cells (**Figure 7I**). In agreement with these results, dramatically enhanced infiltration of activated macrophages, neutrophils and T-cells was observed in the lungs of hACE2 mice and hACE2; *Irgm1^−/−^ Ifnar1^−/−^* mice (**Figure 7J-7O**) than hACE2; *Irgm1^−/−^* mice. Taken together, the data suggest that during SARS-CoV-2 invasion, type 1 IFN response is the major contributor to the protective phenotypes observed in *Irgm1^−/−^* mice.

**Figure 7.**
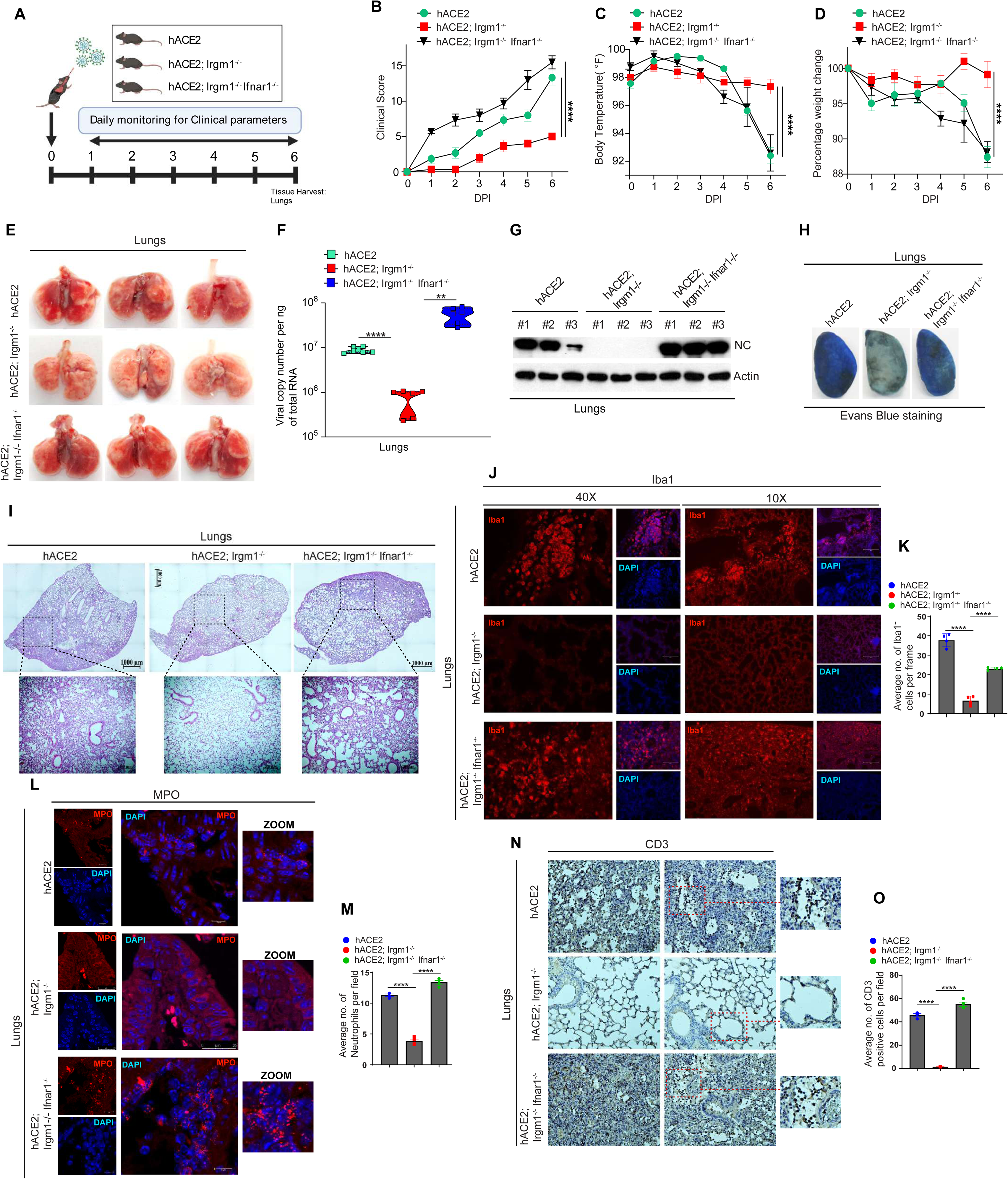
The Irgm1^−/−^ Ifnar1^−/−^ double knockout mice were not protected from SARS-CoV-2 infection and displayed enhanced immunopathology. **A.** Schematic representation of the experimental design depicting timeline of SARS-CoV-2 infection in hACE2, hACE2; *Irgm1^−/−^* and hACE2; *Irgm1^−/−^ Ifnar1^−/−^* transgenic mice. Mice were intranasally inoculated with 5X10^4^ PFU (low dose) of SARS-CoV-2. Image created using Biorender.com. **B-D.** Graph showing **(B)** total clinical scores, **(C)** core body temperature (in °F) and **(D)** percentage change in body weight of hACE2, hACE2; *Irgm1^−/−^* and hACE2; *Irgm1^−/−^ Ifnar1^−/−^* mice infected with SARS-CoV-2 for 6 days. Mean ± SEM, ****p < 0.0001, Two-way ANOVA (Sidak’s multiple comparison test). **E.** Lung images showing gross pathology in hACE2, hACE2; *Irgm1^−/−^* and hACE2; *Irgm1^−/−^ Ifnar1^−/−^* mice infected with SARS-CoV-2 at 6 dpi. **F.** Viral copy number analysis in total RNA isolated from lungs of SARS-CoV-2 infected hACE2, hACE2; *Irgm1^−/−^* and hACE2; *Irgm1^−/−^ Ifnar1^−/−^* mice at 6 dpi (n=6). Mean ± SEM, **p < 0.01, ****p < 0.0001, One-way ANOVA (Tukey’s multiple comparison test). **G.** Western blot analysis of NC protein expression in lung lysates of SARS-CoV-2 infected hACE2, *hACE2; Irgm1^−/−^* and hACE2; *Irgm1^−/−^ Ifnar1^−/−^* mice at 6 dpi. **H.** Evan’s blue staining in lungs of hACE2, hACE2; *Irgm1^−/−^* and hACE2; *Irgm1^−/−^ Ifnar1^−/−^* mice infected with SARS-CoV-2 at 6 dpi. **I.** Representative hematoxylin & eosin stained lung images of SARS-CoV-2 infected hACE2, hACE2; *Irgm1*^−/−^ and hACE2; *Irgm1*^−/−^ *Ifnar1*^−/−^ mice at 6 dpi. Panel below showing zoomed image of lung regions marked in inset. **J-K. (J)** Representative images of IBA1 immunostained lungs sections of SARS-CoV-2 infected hACE2, hACE2; *Irgm1*^−/−^ and hACE2; *Irgm1*^−/−^ *Ifnar1*^−/−^ mice (6 dpi). Images captured at 40X and 10X magnifications. **(K)** Bar graph showing average number of IBA1 positive cells per frame (n=4, mean ± SEM, ****p < 0.0001, One-way ANOVA (Tukey’s multiple comparison test). **L-M. (L)** Representative confocal images of MPO (neutrophils) immunostained lungs of SARS-CoV-2 infected hACE2, hACE2; *Irgm1*^−/−^ and hACE2; *Irgm1*^−/−^ *Ifnar1*^−/−^ mice (6 dpi). Right panels showing zoomed image. **(M)** Bar graph showing average number of MPO positive cells per field (n=4, mean ± SEM, ****p < 0.0001, One-way ANOVA (Tukey’s multiple comparison test). **N-O. (N)** Representative immunohistochemistry images of CD3 (T-cells) immunostained lungs of SARS-CoV-2 infected hACE2, hACE2; *Irgm1*^−/−^ and hACE2; *Irgm1*^−/−^ *Ifnar1*^−/−^ mice (6 dpi). **(O)** Bar graph showing average number of CD3 positive cells per field. n=4, mean ± SEM, ****p < 0.0001, One-way ANOVA (Tukey’s multiple comparison test).

## Discussion

The goal of this study is to define the role of IFN-I in SARS-CoV-2 infection. We used multiple genetically manipulated mouse models and chemical treatments to understand the interaction between the IFN-I system and SARS-CoV-2. Our results show that the IFN-I system is indispensable for protecting the mice from SARS-CoV-2 infection, inflammation, neuroinvasion, immunopathology, and subsequent lethality.

The *Irgm1* knockout mice displayed very mild clinical symptoms with none of the mice undergoing hypothermic conditions. A condition that was shown to be associated with enhanced mortality rates in humans (Fatteh et al, 2021; Maait et al, 2021) The viral load remained low in *Irgm1^−/−^* mice throughout the experiments performed for different time points. The pathological changes in *Irgm1* knockout mice were minor as compared to parental hACE2 mice, which showed severe lung consolidation and fibrosis with enhanced migration of neutrophils, T-Cells, and macrophages. Consistently, the *Irgm1* knockout mice were protected from the lethal cytokine storm observed in the case of hACE2 mice (Arce & Costoya, 2021; Golden et al, 2020; Oladunni et al, 2020; Winkler et al, 2020; Yinda et al, 2021). To be sure that IFN-I is a major contributor to the protection observed in *Irgm1* knockout mice, we generated *Irgm1* and *Ifnar1* double knockout mice. Like hACE2 mice, we found that hACE2; *Irgm1^−/−^ Ifnar1^−/−^* mice were equally susceptible to SARS-CoV-2 infection and displayed similar trend of immunopathology suggesting that IFN-I plays a major role in protecting *Irgm1* knockout mice from SARS-CoV2 invasion. Thus, the preexisting heightened IFN-I immunity in *Irgm1^−/−^* mice could limit the SARS-CoV-2 replication and disease progression. Also, we found that this immunity in hACE2; *Irgm1^−/−^* mice resisted the SARS-CoV-2 infection better than prophylactic IFN-α treatment in hACE2 mice. The development of therapeutics that could boost the host anti-SARS-CoV2 response could increase our preparedness for future viral pandemics especially those related to the β-coronavirus family. In this context, the current and previous studies (Nath et al, 2021) from our lab strongly establish that *Irgm1* (or human IRGM) could be an excellent target for developing host-based broad antiviral therapeutics. Due to the linkage between IRGM loss-of-function mutations and autoimmune diseases, there appears to be a caveat to this hypothesis that such therapeutics may increase the likelihood of autoimmune responses. However, we believe that because these therapeutics would be administered only for a short period of time (as prophylactic measures) during pandemics, epidemics, and endemics, the chance of triggering an autoimmune response will be low, and the benefits may outweigh the risks.

Several recent population-based studies have suggested the effectiveness of the IFN-I system in protecting against a wide range of viruses including SARS-CoV-2 (Abolhassani et al, 2022; Bastard et al, 2022; Duncan et al, 2022; Khanmohammadi et al, 2022; Meyts, 2022; Zhang et al, 2022). Also, studies found that COVID-19 patients with antibodies to IFN-I are vulnerable to severe disease outcomes (Bastard et al, 2021; Bastard et al, 2020; Frasca et al, 2022). In agreement with these studies, we found that hACE2; *Ifnar1^−/−^* mice were highly susceptible to SARS-CoV-2 infection in the lungs and brain. Upon SARS-CoV-2 infection, in these mice, although IFN-I was suppressed, other inflammatory responses such as neutrophil/leukocyte migration, IL-17 signaling, interferon-gamma response, neutrophil degranulation, IL-β, and IL-6 response, TNF-α response, fibrin/clot formation, and fibrosis were hyperinduced. In addition, we found hACE2; *Ifnar1^−/−^* mice were highly susceptible to SARS-CoV-2 neuroinvasion of the brain resulting in microgliosis/astrogliosis, immune cell infiltration, enhanced chemokine/interleukin response and neuropathology’s. Thus, this model could help us to understand the molecular mechanisms driving severe viral diseases in patients with a loss-of-function mutation in IFNAR alleles. Several case reports and patient studies suggest that SARS-CoV-2 can invade the central nervous system (Bauer et al, 2022; Douaud et al, 2022; Jacob et al, 2020; Meinhardt et al, 2021; Rutkai et al, 2022; Seehusen et al, 2022; Sepehrinezhad et al, 2021; Song et al, 2021; Veleri, 2022). Given the hypersusceptibility of hACE2; *Ifnar1^−/−^* mice brain to SARS-CoV-2 infection and surprisingly high probability of loss-of-function mutation in IFNAR1/2 gene in the population, we speculate whether SARS-CoV-2 neuroinvasion is more common in the population with null IFNAR1/2 alleles leading to severe disease outcomes. Taken together, in this study, we have not only defined the role of IFN-I response in SARS-CoV-2 infection but also generated mouse models that could be useful in determining the molecular basis of the heterogeneity in COVID-19 disease outcomes in patients. This knowledge could be very useful for preparedness for future viral pandemics such as COVID-19.

## Materials and Methods

### Ethics statement and biosafety

All animal experiments were approved by the Institutional Animal Ethics Committee (IAEC) of the Institute of Life Sciences, Bhubaneswar, India. All procedures with infectious SARS-CoV-2 were performed in approved high containment BSL3 and A-BSL3 facilities at the Institute of Life Sciences using appropriate and necessary precautionary measures. All SARS-CoV-2-related work was performed with approved standard operating procedures and safety measures were strictly followed.

### Experimental model and subject details

#### Cells and viruses

Vero E6 cells were cultured at 37°C in Dulbecco’s Modified Eagle medium (DMEM; Gibco #10569044) supplemented with penicillin/streptomycin (10,000 units/mL) and 10% fetal bovine serum (FBS). SARS-CoV-2 strain of clade 19A was isolated, characterized, and propagated as described previously (Kumar et al, 2021; Singh et al, 2022; Suresh et al, 2021) (Genbank Accession MW559533). The SARS-CoV-2 Delta strain (GISAID ID EPI_ISL_2775201; hCoV-19/India/TG-CCMB-CIA4413/2021) was isolated, characterized, and propagated as described previously (Tandel et al, 2022).

#### Chemicals

H-151 (STING Inhibitor InvivoGen #inh-h151), GSK583 (RIPK2 inhibitor-Cayman Chemical Company #19739), IFN-α (PBL Assay Science #11200-1), Evans Blue dye (Sigma #E2129), Picrosirius Red Stain Kit (Abcam #ab150681), Solvent Blue 38 (Sigma #S3382), Cresyl violet acetate (SRL #21442), Hematoxylin (Merck #1051750500), Eosin (Merck #1098441000).

### Intranasal inoculation of SARS-CoV-2 in mice

B6.Cg-Tg (K18-ACE2) 2Prlmn/J, commonly referred to as hACE2 mice (JAX stock #034860), Ifnar1 knockout mice, B6(Cg)-Ifnar1 tm1.2Ees/J (JAX stock #028288) mice were purchased from The Jackson Laboratory. Irgm1 knockout (C57BL/6) mice were kindly provided by Dr. Gregory Taylor and maintained as described previously (Bafica et al, 2007; Liu et al, 2013; Mehto et al, 2019; Nath et al, 2021). First, by crossings between hACE2 and *Irgm1^−/−^* mice, we obtained hACE2 (Homozygous) expressing *Irgm1^−/−^* mice. These mice were crossed with *Irgm1^−/−^* mice to obtain hACE2 (hemizygous) expressing *Irgm1* knockout mice (hACE2 (hemizygous); *Irgm1^−/−^*) that were used in experiments. Likewise, hACE2; *Ifnar1* knockout mice (hACE2 (hemizygous); *Ifnar1^−/−^*) were generated by crossing hACE2 (Homozygous); *Ifnar1^−/−^* mice with *Ifnar1^−/−^* mice. hACE2 expressing *Irgm1^−/−^* and *Ifnar1^−/−^* double knockout mice (hACE2 (hemizygous); *Irgm1^−/−^ Ifnar1^−/−^*) were generated by crossing hACE2 (Homozygous); *Irgm1^−/−^ Ifnar1^−/−^* mice with *Irgm1^−/−^ Ifnar1^−/−^* mice. All mice strains were subsequently bred and housed on a 12/12 h light/dark cycle with food pellets and water available *ad libitum*.

8-10 weeks old male and female mice were used in the present study. For each experiment, age-matched littermates were used. Intranasal delivery of SARS-CoV-2 (low dose, 5 x 10^4^ plaque-forming units (pfu) per mouse; high dose, 5 x 10^5^ pfu per mouse) or delta variant (5 x 10^4^ pfu per mouse) was performed under anesthetized animals using a mixture of ketamine (50mg/kg) and xylazine (5mg/kg) injected intraperitoneally. Mice were monitored daily for scoring clinical symptoms, body weight, and core body temperature until days as indicated. For the survival curve experiment, the mice were given SARS-CoV-2 inoculum intranasally and monitored daily for the clinical symptoms, body weight, and core body temperature until 15 dpi. The mice were sacrificed on different days post-infection and various organs were harvested and processed for respective experiments. All efforts were made to minimize animal suffering. Gross lung images were captured with Canon IXUS 185 camera for the presence of lesions.

### Clinical scoring

Mice were monitored daily for scoring clinical symptoms, changes in core body temperature, and body weight post-viral infection. The clinical scores were assigned as follows: normal (without any symptoms), 0; Shivering occasionally, 1; Shivering constantly, 2; Hunchback while sitting, 3; Hunchback while sitting and walking, 4; Abnormal rapid breathing, 4; Slow labored breathing, 5; Slow when stimulated, 5; No response when stimulated, 6 (euthanasia stage).

### Evan’s Blue experiment

hACE2, hACE2; *Irgm1^−/−^*, and hACE2; *Irgm1^−/−^ Ifnar1^−/−^* mice were infected with SARS-CoV-2 and administered with Evan’s Blue dye (1%) intraperitoneally on the 5th day post-infection and sacrificed after 24 h. Mice were transcardially perfused with 1X PBS (pH 7.4) followed by gross image capture of the lungs.

### Inhibitors and IFN-α treatment in mice

hACE2; *Irgm1^−/−^* mice were intraperitoneally injected with GSK583 (25mg/kg body weight) in PBS containing 10% DMSO, 40% PEG 300, and 5% Tween-80 or H-151 (750 nmol/mouse) in PBS containing 10% Tween-80 or IFN-α (2X10^4^ U/mouse) in PBS containing 0.1% BSA daily for 2 days. On the day of infection, mice were injected with GSK583 or H-151 or IFN-α, 4 h prior to SARS-CoV-2 infection. Similarly, control mice were administered with vehicle only followed by infection. All mice were routinely monitored for scoring of clinical symptoms.

### Isolation of mice bone marrow cells and differentiation into macrophages

Isolation and differentiation of mice bone marrow cells into macrophages are performed as described previously (Mehto et al, 2019; Nath et al, 2021). Briefly, mice were sacrificed and bone marrow cells from the tibia and femur were flushed out in RPMI medium. Further, RBC lysis buffer containing (155 mM NH_4_Cl, 12 mM NaHCO_3,_ and 0.1 mM EDTA) was used to remove RBCs. BMDMs were grown in RPMI-1640 media (Gibco #61870127) supplemented with penicillin/streptomycin (10,000 units/ml), 5 mM L-glutamine, glucose (5%), sodium pyruvate (1 mM), HEPES buffer, 10% FBS, containing 20 ng/ml mouse M-CSF (Gibco #PMC2044) for 5 days. The media was replacement every alternate day with fresh media containing M-CSF.

### Transient transfection with siRNA

The freshly isolated bone marrow cells were electroporated (Neon, Invitrogen #MPK5000; setting: 1400 V, 10 ms, 3 pulses using 100 µl tip #MPK10096) with non-specific siRNA (30 nM) or specific siRNA (30 nM) and were differentiated and incubated for 48 h in RPMI medium (10% FBS) containing 20 ng/mL mouse M-CSF. A second round of siRNA transfection was performed with Lipofectamine RNAiMAX transfection reagent (Invitrogen #13778075) after 48 h and incubated for 24 h in the fresh media containing M-CSF. The next day, transiently transfected cells were infected with the SARS-CoV-2 variant (MOI = 0.5) for 16 h. For RNA extraction, the cells were collected in TRIzol (Invitrogen #15596018). The following siRNAs was used in the present study: Non-targeting siRNA (SMARTpool: siGENOME ns siRNA; Dharmacon #D-001206-13-20), Mouse ISG15 siRNA (SASI_MM02_00357360), RSAD2 siRNA (SASI_MM01_00031984), IFITM3 siRNA (SASI_MM01_00156543), MX1 siRNA (SASI_MM01_00166144), OAS1A siRNA (SASI_MM01_00197487).

### Quantification of viral copy number by digital droplet PCR (ddPCR) method

Total RNA was isolated from the lungs of SARS-CoV-2 infected hACE2 mice and cDNA synthesis was carried out from 1 µg of RNA. The cDNA sample was serially diluted every 10-folds with nuclease-free water and subjected to quantification by ddPCR method as per the manufacturer’s protocol. Briefly, each dilution was mixed with the viral Nucleocapsid primer set and droplet generation was performed into the DG8 Droplet Generator Cartridge for each serial dilution in the QX200 Droplet Generator (Bio-Rad). Following droplet generation, the droplets were transferred carefully into a 96-well PCR plate and were sealed with foil using a PX1 Plate Sealer. PCR amplification of the droplets was performed using C1000 Touch Thermal cycler (Bio-Rad). Individual droplets were then read in QX200 Droplet Reader (Bio-Rad) and analyzed using QuantaSoft Software to obtain absolute viral copy numbers. The same serial dilutions were used for qRT-PCR with SARS-CoV-2 nucleocapsid-specific primers to obtain Ct values corresponding to viral copy number. Based on these calculations, the Ct values obtained from the qRT-PCR with SARS-CoV-2 nucleocapsid-specific primers in tissue samples were converted to viral copy numbers.

### Western blotting

Tissue lysate preparation and immunoblotting analysis were performed as described previously (Mehto et al, 2019). Briefly, mice tissue samples were subjected to homogenization in the radio-immunoprecipitation assay (RIPA) buffer (1 mM, EDTA; 20 mM Tris, pH 8.0; 0.1% Sodium deoxycholate; 0.5 mM, EGTA; 1% IGEPAL; 10% glycerol; 150 mM NaCl) along with phosstop, 1 mM PMSF and protease inhibitor cocktail using a glass dounce homogenizer. Following centrifugation at 4°C, the lysates were separated via SDS-PAGE and transferred onto a nitrocellulose membrane (Bio-Rad). The membrane was subjected to blocking for 1 h in 5% skimmed milk followed by incubation in primary antibody overnight at 4°C. The membrane was then washed thrice with 1X PBS/PBST and incubated for 1 h with HRP conjugated secondary antibody. After washing with PBS/PBST the membrane was developed using the enhanced chemiluminescence reagent (Thermo Fisher #32132X3).

### Antibodies

Primary antibodies used in Western blotting with dilutions: Actin (Abcam #ab6276; 1:5000), Irgm1 (CST #14979; 1:1000), SARS-CoV-2 nucleocapsid (Invitrogen #PA5-81794; 1:1000), Spike (CST #56996S; 1:1000), ACE2 (R&D systems #AF933; 1:1000). HRP conjugated secondary antibodies were purchased from Promega (1:5000) or Novus (1:5000).

Primary antibodies used in immunohistochemical assays with dilutions: SARS-CoV-2 nucleocapsid (Invitrogen #PA5-81794; 1:1000), CD3ɛ (CST #99940; 1:200), MPO (Invitrogen #PA5-16672; 1:200), IBA1 (Abcam #ab178846; 1:1000), MBP (CST #78896S, 1:1000), EEA1 (CST #3288; 1:100), Collagen I (Invitrogen #PA1-26204; 1:150), GFAP (CST #3670; 1:200).

### Lungs pathological examination by hematoxylin and eosin (H and E) staining

The lung tissue sections were deparaffinized in xylene and hydrated through an ethanol gradient followed by 1X PBS (pH 7.4) wash. For H and E staining, the sections were stained with hematoxylin for 5 min followed by washing in water to remove excess stain. The sections were then incubated in Scott’s tap water followed by eosin staining. The sections were rinsed, dehydrated in absolute ethanol, cleared in xylene, and mounted with permanent media (VectaMount #H-5000). The slides were observed under ZEISS ApoTome.2 microscope.

### Picrosirius red staining

The lung tissue sections were subjected to deparaffinization in xylene and hydrated through an ethanol gradient followed by distilled water wash. Adequate picrosirius red staining solution was applied to completely cover the lung tissue sections and was incubated for 1 h. The slides were quickly rinsed twice in acetic acid solution followed by rinsing in absolute alcohol. The sections were dehydrated twice in absolute alcohol. The slides were cleared in xylene and mounted with permanent media (VectaMount #H-5000). The slides were observed under ZEISS ApoTome.2 microscope.

### Luxol fast blue staining

The brain sections were deparaffinized in xylene and hydrated to 95% ethanol followed by incubation in luxol fast blue solution (0.1%) at 56°C overnight. The next day, the excess stain was rinsed off with 95% ethanol followed by distilled water. The sections were then differentiated in lithium carbonate solution (0.05%) for 15 seconds followed by rinsing in distilled water and counterstained with cresyl violet solution (0.1%) for 1 minute. The slides were dehydrated through 95% ethanol followed by absolute ethanol. Finally, the sections were cleared in xylene and mounted with permanent media (VectaMount #H-5000). The slides were visualized under ZEISS ApoTome.2 microscope.

### Immunohistochemistry

For immunohistochemical staining, lungs and brain tissue sections were deparaffinized in xylene and hydrated through an ethanol gradient followed by 1X PBS (pH 7.4) wash. Endogenous peroxidase activity was blocked at room temperature by H_2_O_2_ as per required. Antigen retrieval was carried out in 10mM sodium citrate buffer (pH 6.0) for 10 min followed by permeabilization with 1X PBS (pH 7.4) containing 0.25% Triton X-100, blocked with normal goat serum. The sections were incubated overnight at 4°C with the primary antibody. Next day, the sections were washed with 1X PBS (pH 7.4) containing 0.05% Tween 20, followed by incubation with goat anti-rabbit IgG (H+L) or goat anti-mouse IgG (H+L) either HRP or Alexa Fluor 568 or Alexa Fluor 488 conjugated secondary antibody for 1 h at room temperature. Sections were then washed and developed with DAB peroxidase substrate kit (VectorLabs, #SK-4100). Sections were finally counterstained with hematoxylin stain followed by dehydration through an alcohol gradient. Sections were cleared in xylene and mounted with permanent media (VectaMount #H-5000). For immunofluorescence, the sections were washed and counterstained with DAPI for 1 min followed by incubation with autofluorescence quencher (Vector Labs #SP-8400). Sections were finally mounted with Vectashield Vibrance Antifade mounting media. The slides were visualized under ZEISS ApoTome.2 microscope or Leica TCS SP8 STED confocal microscope.

The immunohistochemical data were quantified by counting the number of positive cells per frame (n=4 mice per group). 10 fields per section per mouse were randomly captured and quantified.

### RNA isolation and quantitative real-time PCR

The total RNA extraction from tissues was performed using TRIzol^TM^ (Catalogue# 15596026)by following the manufacturer’s procedure. cDNA synthesis was carried out using the high-capacity DNA reverse transcription kit (Applied Biosystems #4368813) and qRT-PCR was performed using Power SYBR green PCR master mix (Applied Biosystems, #4367659) as per the manufacturer’s procedure. For assay normalization, the housekeeping gene β-Actin was used. The fold-change in expression was calculated by the 2^−ΔΔCt^ method.

The following gene-specific primer sequence for cytokines, chemokines, and ISGs was used for qRT PCR:

#### qRT primers

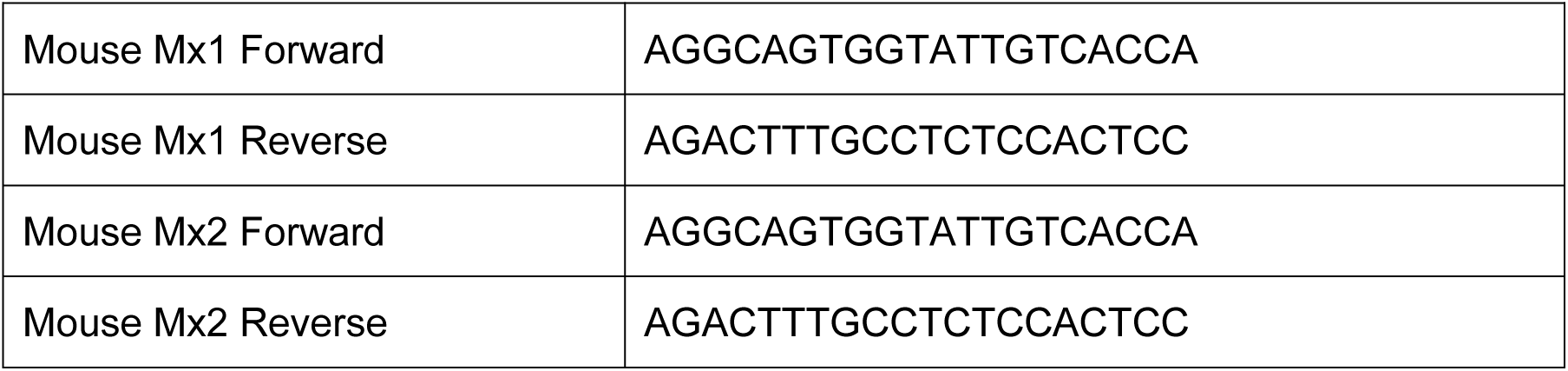
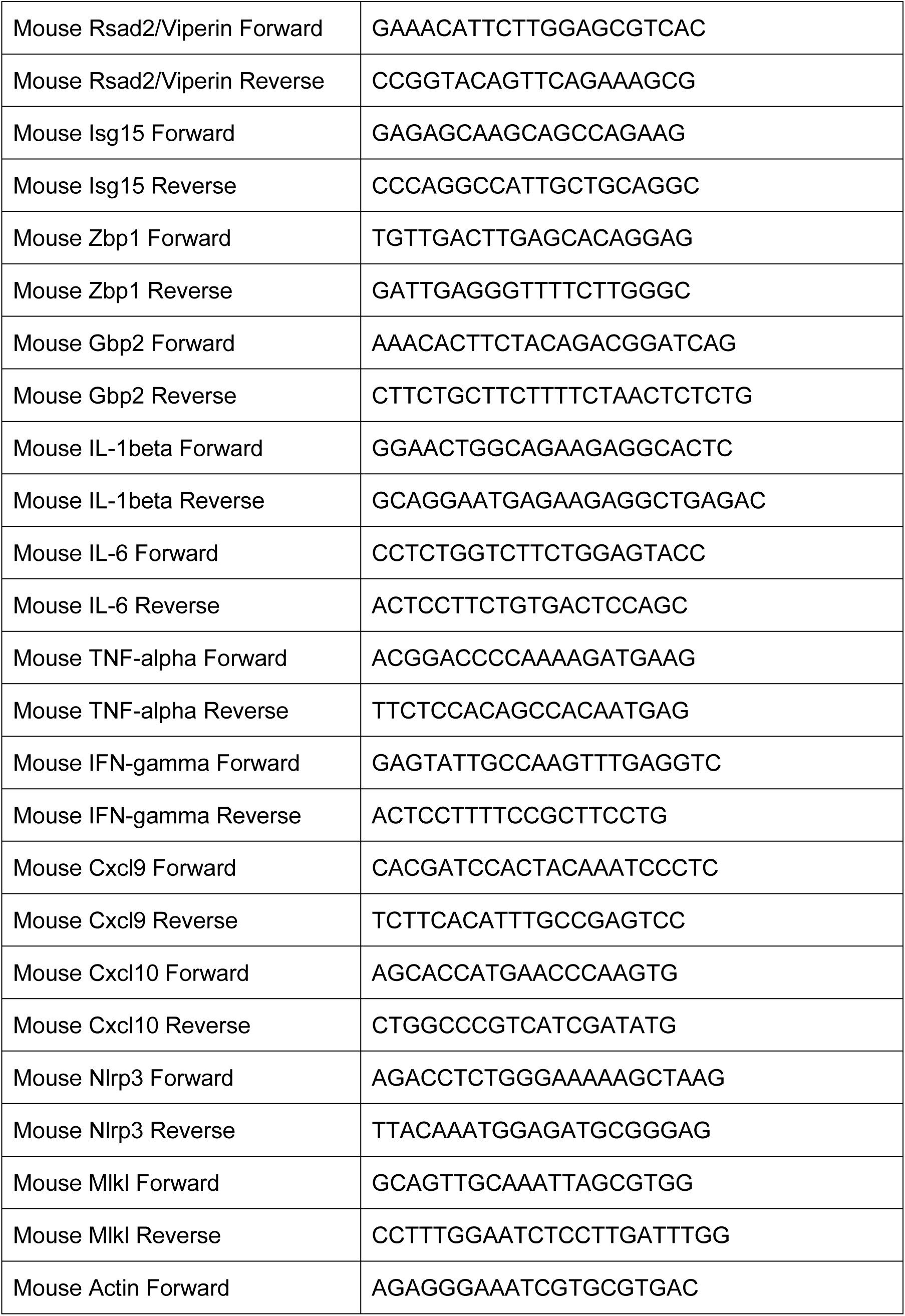
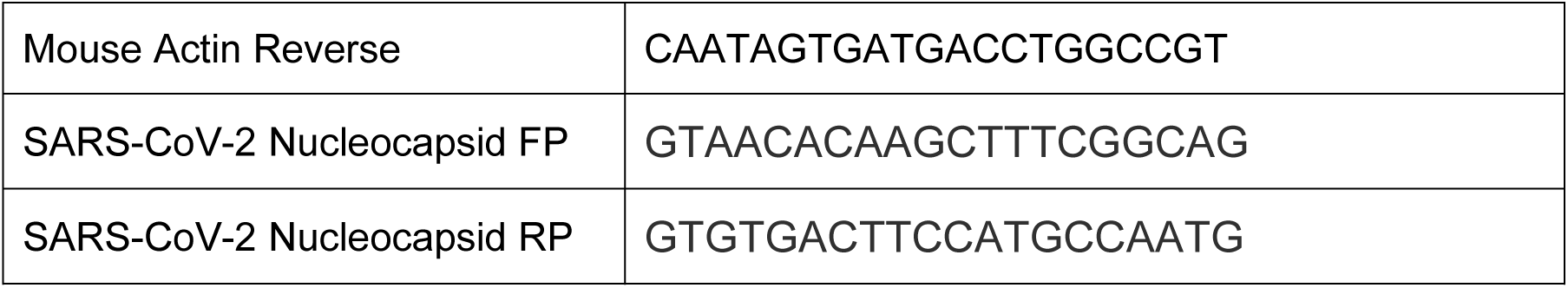

### RNA sequencing sample preparation

The total RNA was extracted from uninfected and SARS-CoV-2 infected mice lung tissue using RNeasy mini kit (QIAGEN #74104). The quantity of RNA was determined using Qubit RNA assay kit with Qubit 4.0 and the quality of RNA was tested using agarose gel electrophoresis and High Sensitivity Tape station Kit (Agilent 2200, #5067-5576, #5067-5577 and #5067-5578). After assessing the quality of RNA, ∼900 ng of total RNA was taken for library preparation using NEBNext^®^Ultra™ II Directional RNA Library kit for Illumina (# E7760L) and NEBNext Poly (A) mRNA Magnetic Isolation Module (# E7490L) as per manufacturer’s protocol. The prepared library was quantified using Qubit dsDNA assay kit (Invitrogen, Q32851) followed by quality check (QC) and fragment size distribution using a High Sensitivity Tape station Kit (Agilent 2200, #5067-5584 and #5067-5585). The library was sequenced using the HiSeq 4000 Illumina platform.

### RNA sequencing data processing and gene expression analysis

The paired-end (PE) reads quality checks for each sample were carried out using FastQC v.0.11.5 (http://www.bioinformatics.babraham.ac.uk/projects/fastqc/). The adapter sequence was trimmed using the BBDuk version 37.58 version 37.58 and the alignment was performed using STAR v.2.5.3a with default parameters(Dobin et al, 2013) with human hg38 genome build, gencode v21 gtf 9GRCh38) from the gencode. The duplicates were discarded using Picard-2.9.4 (https://broadinstitute.github.io/picard/) from the aligned bam files and read counts were generated using featureCount v.1.5.3 from subread-1.5.3 package (https://bioinf.wehi.edu.au/) with Q = 10 for mapping quality. The count files were used as input for downstream differential gene expression analysis with DESeq2 version 1.14.1 9 (Love et al, 2014). The genes with read counts of ≤ 10 in any comparison were discarded followed by count transformation and statistical analysis using DESeq “R”. The “P” value were adjusted using the Benjamini and Hochberg multiple testing correction and the differentially expressed genes were identified (fold change of ≥1.5, P-value < 0.05). A unified non-redundant gene list was made for different comparisons and subjected to gene ontology (GO) analysis using the reactome database (https://reactome.org/). The top pathways (p < 0.05) were used for generating heat maps using Complexheatmap (Version 2.0.0) through unsupervised hierarchical clustering. The expression clusters were annotated based on enriched GO terms. Normalized gene expression was used to generate the boxplots with a median depicting the trends in the expression across the different conditions using ggplot2 [version 3.3.5]. The pathways analysis was performed using Metascape database (https://metascape.org/gp/index.html#/main/step1). The top pathways (p < 0.05) were taken for constructing bubble plots using ggplot2 [version 3.3.5].

### Software and statistical significance

In the present study, GraphPad Prism 9 was used to plot and analyze the data. The number of biological replicates and statistical tests are mentioned in figure legends. Pairwise comparisons between the groups were analyzed by unpaired Student’s t-test. Comparison between more than two groups was done by one-way or two-way ANOVA test with multiple comparisons test as mentioned in figure legends. The p-value < 0.05 was considered statistically significant for all the experiments.

### Graphical illustrations

Graphical illustrations were made with Biorender.com.

## Supporting information

Supplemental data set 1

Supplemental data set 2

Supplemental data set 3

Supplemental data set 4

Supplemental data set 5

Supplemental data set 6

## Acknowledgments

This work is funded by the Wellcome Trust/Department of Biotechnology (DBT) India Alliance (IA/I/15/2/502071) fellowship to Santosh Chauhan. Funding from EMBO Global investigator fellowship to Santosh Chauhan is greatly acknowledged. We gratefully acknowledge the support of the Institute of Life Sciences’ central facilities (BSL3, Animal House, microscopy, FACS, and sequencing) funded by the Department of Biotechnology (India)

## Disclosure and competing interests statement

The authors declare that they have no conflict of interest

## Supplementary Figure legends

**Supplementary Figure 1.**
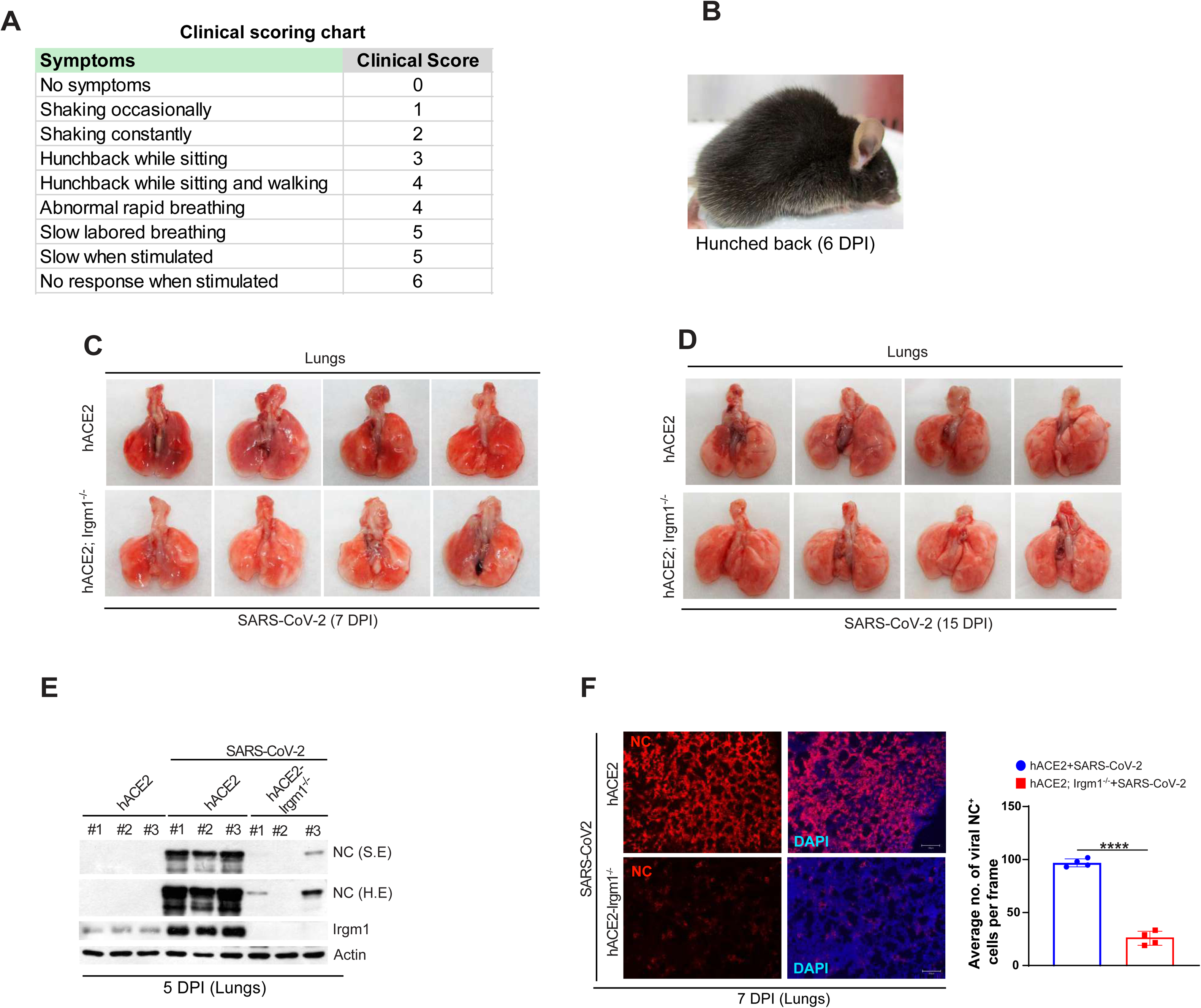
hACE2; Irgm1^−/−^ mice displayed mild clinical scores and pathology. **A.** Table showing symptoms and the corresponding clinical score of the mice infected with SARS-CoV-2. **B.** Representative image of SARS-CoV-2 infected hACE2 mouse with prominent hunched back phenotype at 6 dpi. **C-D.** Lung images showing gross pathology in hACE2 and hACE2; *Irgm1*^−/−^ mice infected with SARS-CoV-2 **(C)** High dose (5X10^5^ PFU) at 7 dpi, and **(D)** Low dose (5X10^4^ PFU) at 15 dpi. **E.** Western blot analysis of NC protein expression in lung lysates of uninfected and SARS-COV-2 infected hACE2, hACE2; *Irgm1*^−/−^ mice at 5 dpi. **F.** Representative images of SARS-CoV-2 nucleocapsid immunostained lungs of SARS-CoV-2 infected hACE2 and hACE2; *Irgm1*^−/−^ mice (7 dpi). Nucleus is stained with DAPI. Right panel showing bar graphs for average number of nucleocapsid positive cells per frame (10 fields from each experiment, n=4 experiments, mean ± SEM, ****p < 0.0001, Student’s unpaired t-test).

**Supplementary Figure 2.**
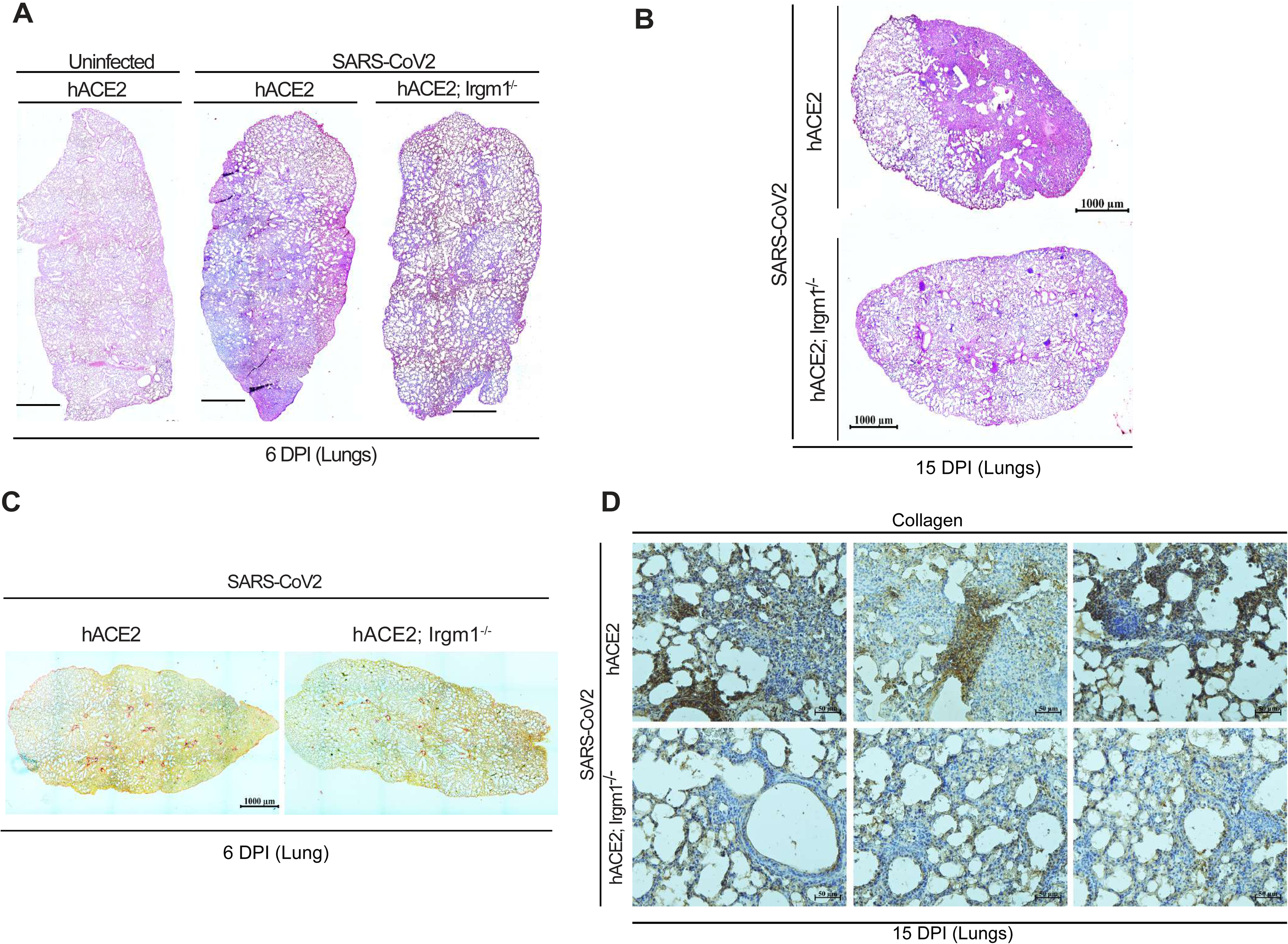
hACE2; Irgm1^−/−^ mice are protected from SARS-CoV-2 infection-mediated immunopathology. **A-B.** Representative hematoxylin & eosin stained lung images of mock and SARS-CoV-2 infected hACE2 and hACE2; *Irgm1*^−/−^ mice at (**A)** 6 dpi and (**B)** 15 dpi. **C.** Representative picrosirius red stained lung images of SARS-CoV-2 infected hACE2 and hACE2; *Irgm1*^−/−^ mice showing collagen deposition (in red) at 6 dpi. **D.** Representative immunohistochemistry images of collagen immunostained lungs of SARS-CoV-2 infected hACE2 and hACE2; *Irgm1*^−/−^ mice (15 dpi).

**Supplementary Figure 3.**
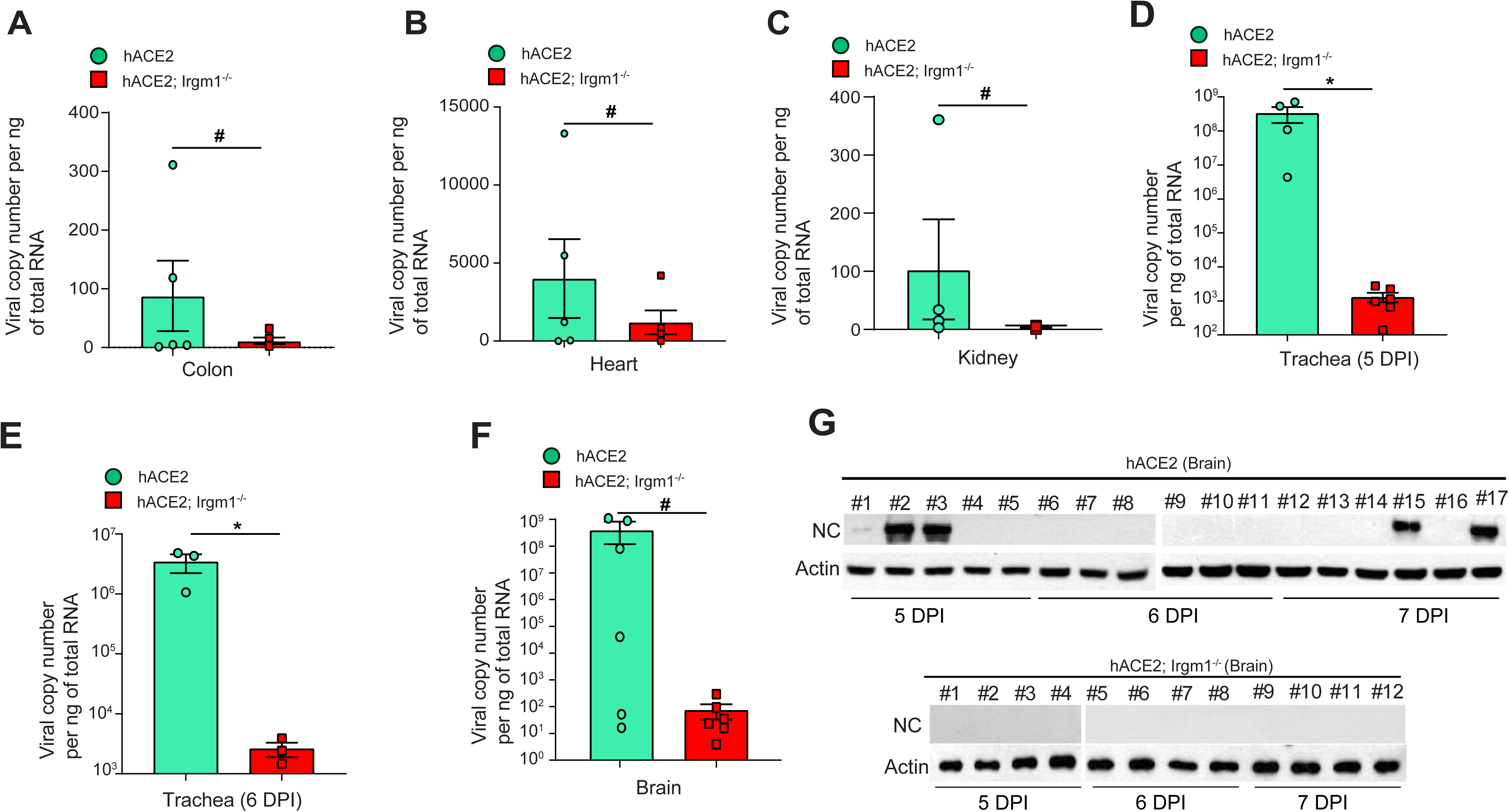
Milder SARS-CoV-2 infection in hACE2; Irgm1^−/−^ mice vital organs. **A-F.** Viral copy number analysis in total RNA isolated from **(A)** colon (6 dpi), **(B)** heart (6 dpi), **(C)** kidney (6 dpi), **(D)** trachea (5dpi), **(E)** trachea (6 dpi), and **(F)** brain (5dpi) of SARS-CoV-2 infected hACE2 and hACE2; *Irgm1*^−/−^ mice. Mean ± SEM, *p < 0.05, ^#^ ns, Student’s unpaired t-test. **G.** Western blot analysis of NC protein expression in brain lysates of SARS-CoV-2 infected hACE2 and hACE2; *Irgm1*^−/−^ mice at 5, 6, and 7 dpi.

**Supplementary Figure 4.**
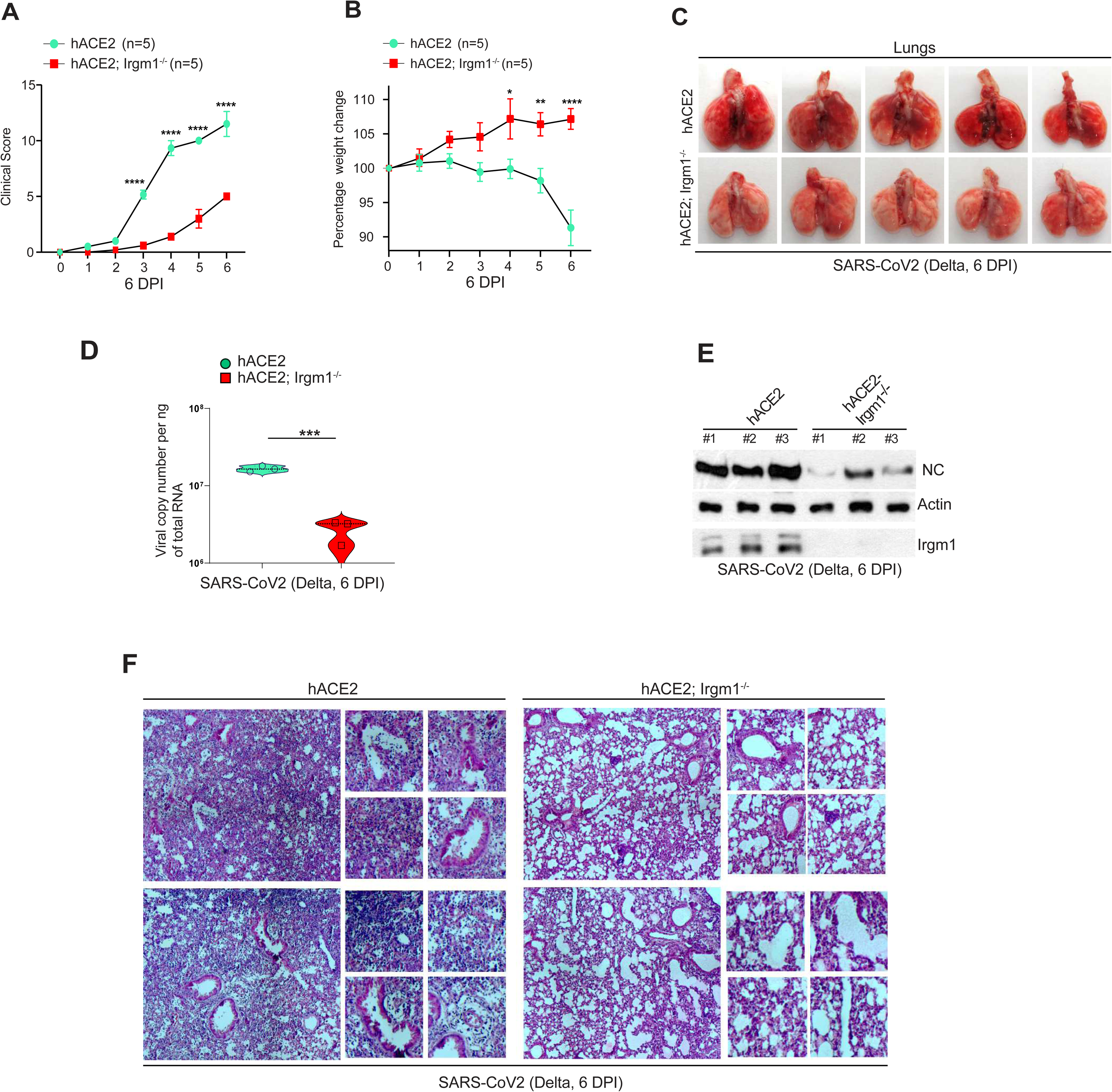
hACE2; Irgm1^−/−^ are resistant to infection of delta variant of SARS-CoV-2. **A-B.** Graphs depict **(A)** clinical scores, and **(B)** percentage change in body weight of SARS-CoV-2 (5X10^4^ PFU) infected hACE2 and hACE2; *Irgm1*^−/−^ mice for 6 days. mean ± SEM, *p < 0.05, **p < 0.01 and ****p < 0.0001, Two-way ANOVA (Sidak’s multiple comparison test). **C.** Lung images showing gross pathology in hACE2 and hACE2; *Irgm1*^−/−^ mice infected with SARS-CoV-2 (5X10^4^ PFU) at 6 dpi. **D.** Viral copy number analysis in total RNA isolated from lungs of SARS-CoV-2 infected hACE2 and hACE2; *Irgm1*^−/−^ mice at 6 dpi. Mean ± SEM, ***p < 0.001, Student’s unpaired t-test. **E.** Western blot analysis of NC protein expression in lung lysates of SARS-CoV-2 infected hACE2 and hACE2; *Irgm1*^−/−^ mice at 6 dpi. **F.** Representative hematoxylin & eosin stained lung images of SARS-CoV-2 infected hACE2 and hACE2; *Irgm1*^−/−^ mice at 6 dpi.

**Supplementary Figure 5.**
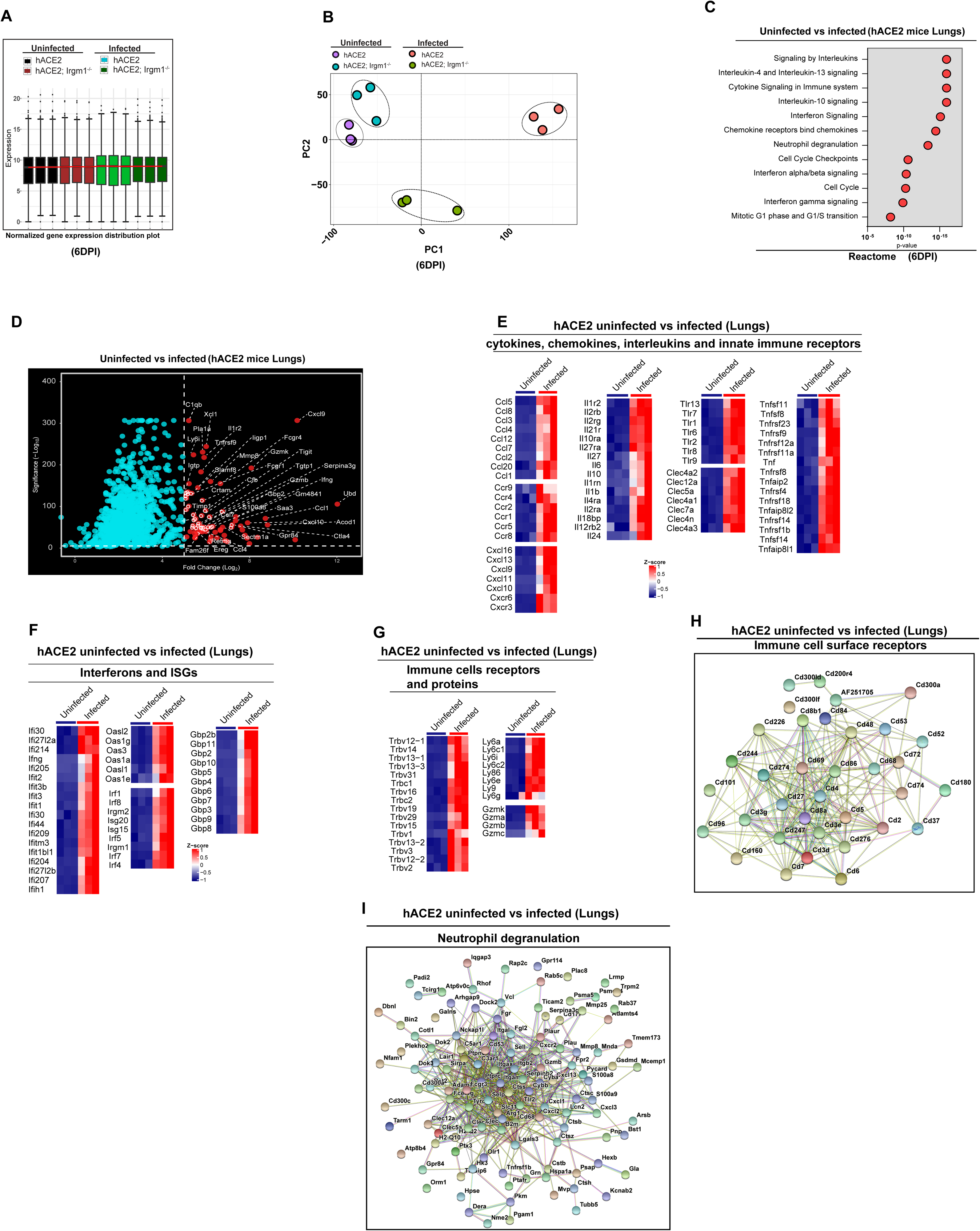
Cytokine storm in lungs of infected hACE2 mice. **A.** Box whisker distribution plot with normalized log expression of the differentially expressed genes (p<0.05, Wald Chi-Squared test, n=3) at 6 dpi **B.** Principal component analysis (PCA) of transcriptomes (p < 0.05, n=3) of different experimental groups, uninfected hACE2, hACE2; *Irgm1*^−/−^ and SARS-CoV-2 infected hACE2, hACE2; *Irgm1*^−/−^ mice lungs at 6 dpi. **C.** Gene ontology-based pathway analysis using Reactome showing enriched pathway in lungs of SARS-CoV-2 infected hACE2 mice (compared to uninfected hACE2; 3 fold, p<0.05, Wald Chi-Squared test, n=3) at 6 dpi. **D.** Volcano plot depicting top upregulated genes in lungs of SARS-CoV-2 infected hACE2 mice (compared to uninfected hACE2; 3 fold, p<0.05, n=3). **E-G.** The heatmaps of genes that are **(E)** cytokines, chemokines, interleukins, innate immune receptors or **(F)** interferons and ISGs or **(G)** immune cells (T-cell, B-cell, macrophages, Neutrophils, NK cells) receptors and proteins, in lungs of SARS-CoV-2 infected hACE2 mice vs uninfected hACE2. (3 fold, p<0.05, Wald Chi-Squared test, n=3). **H-I.** Functional networks (using STRING) of genes related to **(H)** immune cell surface receptors **(I)** Neutrophil degranulation that are induced in lungs of SARS-CoV-2 infected hACE2 compared to uninfected hACE2 mice.

**Supplementary Figure 6.**
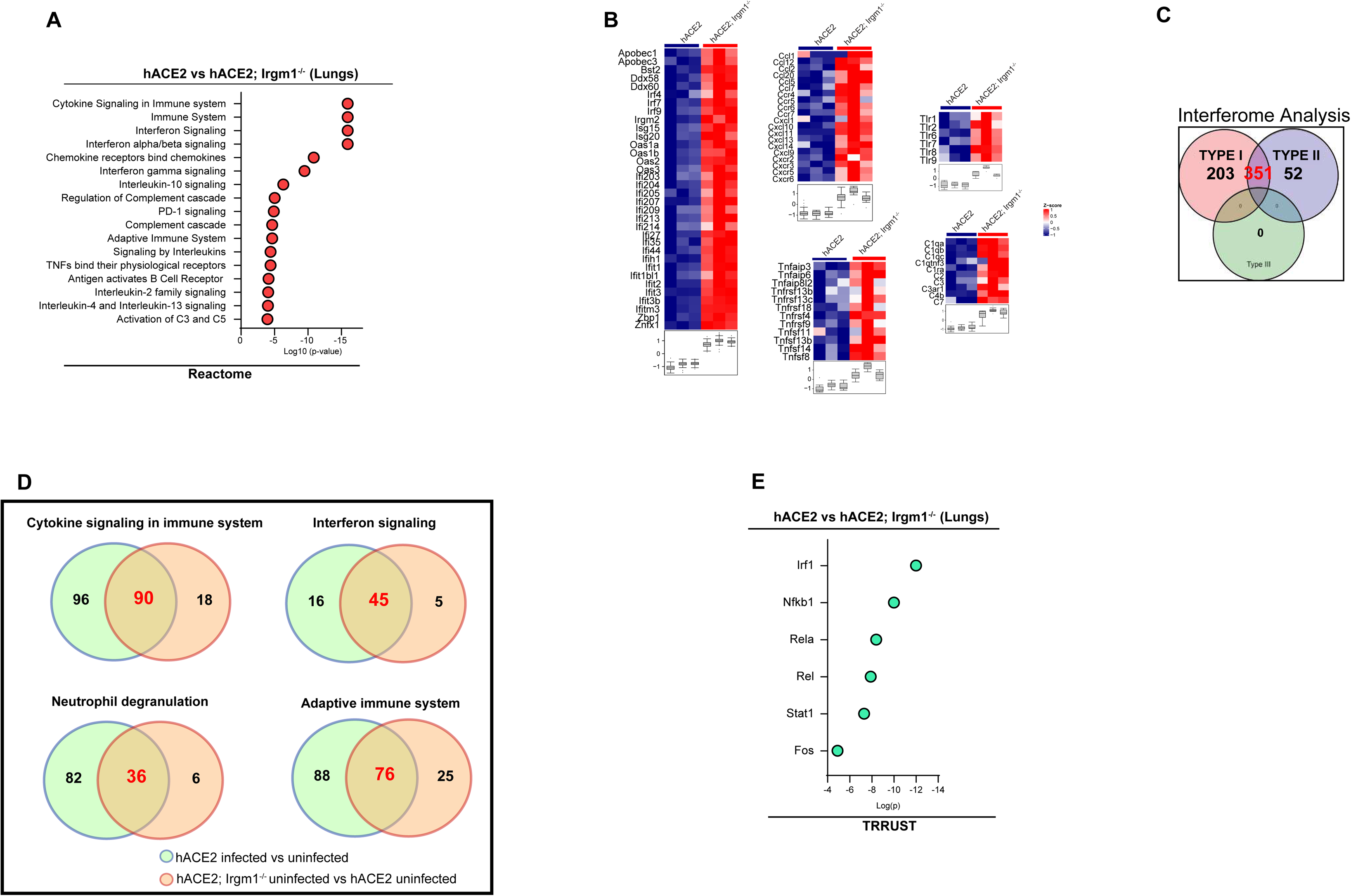
Blunted inflammatory response in Irgm1^−/−^ mice. **A.** Reactome pathway analysis showing enriched pathway in lungs of uninfected hACE2; *Irgm1*^−/−^ mice compared to hACE2 (1.5 fold, p<0.05, Wald Chi-Squared test, n=3). **B.** The heatmap representation of genes related to pathways of interferon signaling, cytokine/chemokines signaling, and complement signaling in lungs of uninfected hACE2; *Irgm1*^−/−^ mice compared to hACE2 mice (1.5 fold, p<0.05, Wald Chi-Squared test, n=3). **C.** Interferome database analysis with a set of genes induced in lungs of uninfected hACE2; *Irgm1*^−/−^ mice compared to hACE2 (1.5 fold, p<0.05, Wald Chi-Squared test, n=3). The Venn diagram depicts the total number of type I, type II, and type III IFN-regulated genes upregulated in lungs of hACE2; *Irgm1*^−/−^ mice compared to hACE2. **D.** The Venn diagrams depicts overlap between genes (related to different pathways) that are upregulated in hACE2; *Irgm1^−/−^* mice lungs in basal conditions (compared to hACE2 mice) and that are upregulated in SARS-CoV-2 infected hACE2 mice lungs (compared to uninfected hACE2 mice lungs). **E.** TRRUST analysis from the transcriptome induced in hACE2; *Irgm1^−/−^* mice lungs in basal conditions compared to hACE2 mice (p<0.05, 1.5 folds, n=3) to predict transcription factors that drive the gene expression.

**Supplementary Figure 7.**
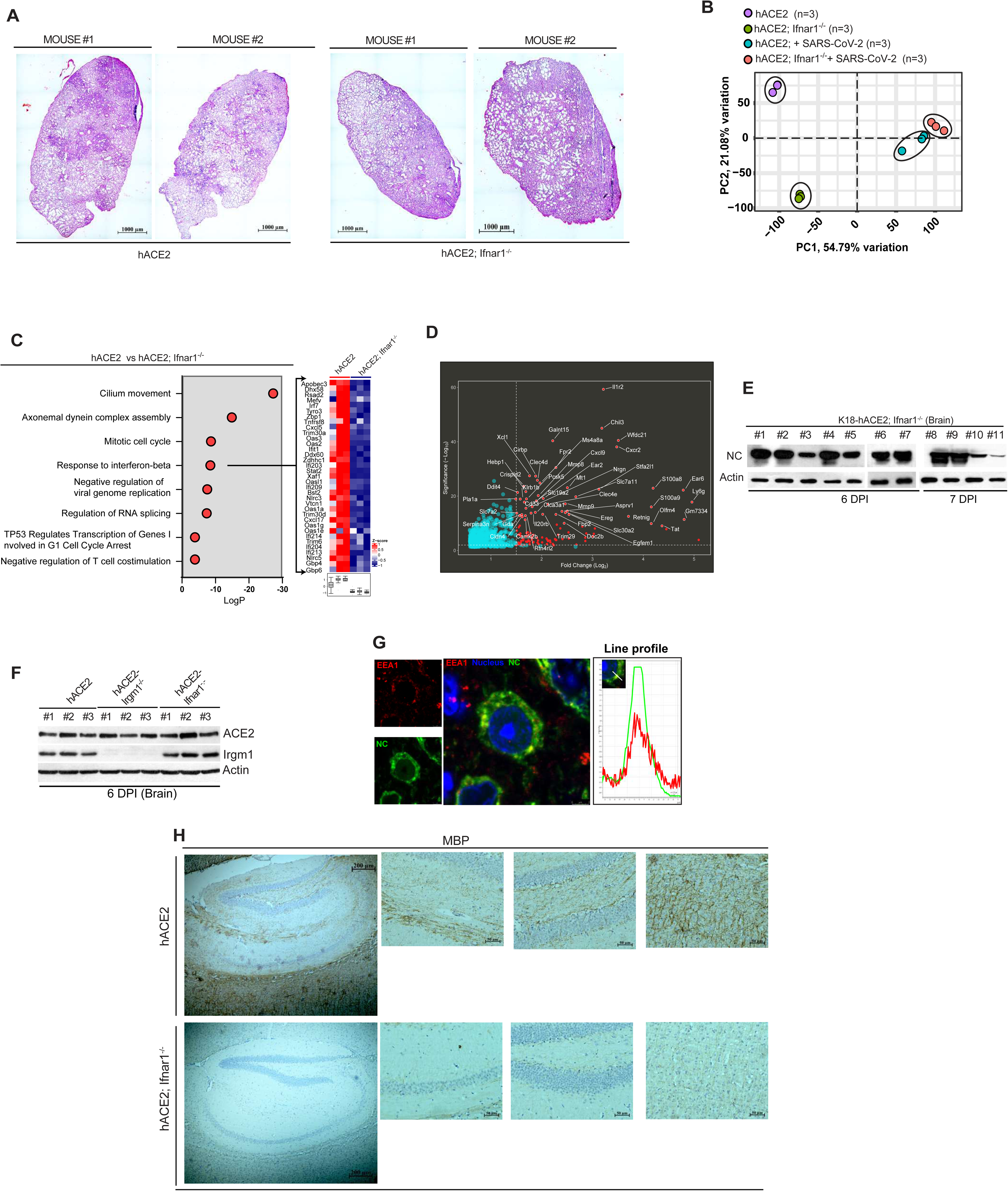
Enhanced SARS-CoV-2 infection, pathology and cytokine storm in hACE2; Ifnar1^−/−^ mice. **A.** Representative hematoxylin & eosin stained lung images of SARS-CoV-2 infected hACE2 and hACE2; *Ifnar1*^−/−^ mice. **B.** Principal component analysis (PCA) of transcriptomes (p < 0.05, n=3) of different experimental groups, uninfected hACE2, hACE2; *Ifnar1*^−/−^ and SARS-CoV-2 infected hACE2, hACE2; *Ifnar1*^−/−^ mice lungs. **C.** Reactome pathway analysis from geneset that is suppressed (>50%, p<0.05, Wald Chi-Squared test, n=3) in hACE2; *Ifnar1*^−/−^ compared to hACE2 mice in basal conditions. Heatmap depicts the genes related to the “response to interferon-beta” pathway. **_D._** Volcano plot depicting top 50 upregulated genes in lungs of SARS-CoV-2 infected hACE2; *Ifnar1*^−/−^ compared to hACE2 mice (1.5 fold, p<0.05, n=3) at 7 dpi. **E.** Western blot analysis of NC protein expression in brain lysates of SARS-CoV-2 infected hACE2; *Ifnar1*^−/−^ mice at 6 and 7 dpi. **F.** Western blot analysis of ACE2 protein expression in brain lysates of SARS-CoV-2 infected hACE2, hACE2; *Irgm1*^−/−^ and hACE2; *Ifnar1*^−/−^ mice at 6 dpi. **G.** Representative confocal images of brain section of SARS-CoV-2 infected hACE2; *Ifnar1*^−/−^ mice at immunostained with EEA1 (red) and viral nucleocapsid (NC, green). Line profile: colocalization analysis using line intensity profiles. Scale bar: 3 μm. Zoom panel is digital magnification. Nucleus stained with DAPI. **H.** Representative immunohistochemistry images of MBP immunostained brain sections of SARS-CoV-2 infected hACE2 and hACE2; *Ifnar1*^−/−^ mice (7 dpi). Right panels showing zoomed images.

**Supplementary Figure 8.**
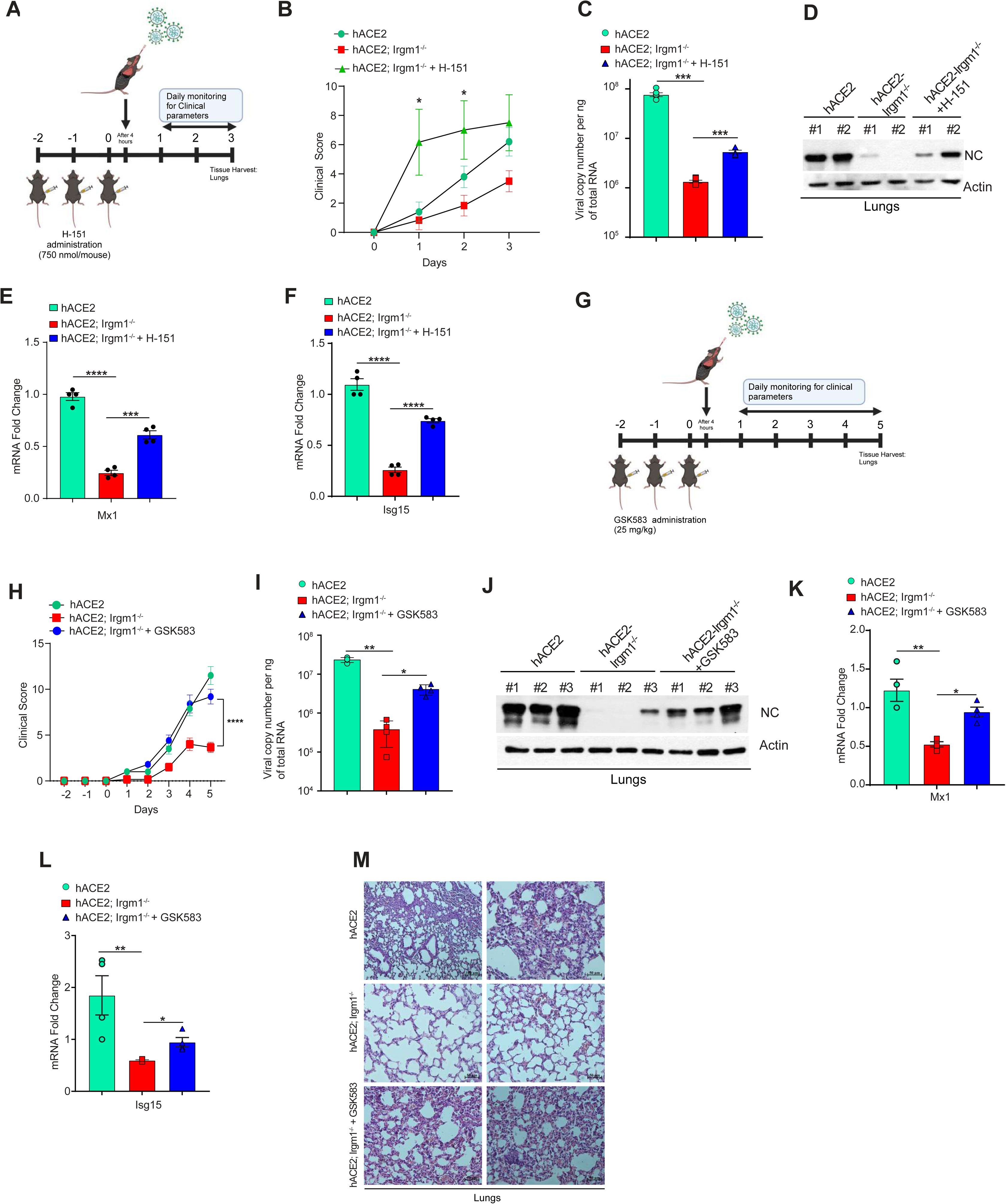
STING and RIPK2 inhibition leads to enhanced SARS-CoV-2 infection in hACE2; *Irgm1*^−/−^ mice. **A.** Schematic representation of the experimental design depicting H-151 treatment (750nmol/mouse) followed by SARS-CoV-2 (5X10^4^ PFU) infection. Image created using Biorender.com. **B.** Graph showing total clinical scores of hACE2, hACE2; *Irgm1*^−/−^ and hACE2; *Irgm1*^−/−^ + H-151 treated mice infected with SARS-CoV-2 for 3 days. Mean ± SEM, *p < 0.05, Two-way ANOVA (Sidak’s multiple comparison test). **C.** Viral copy number analysis in total RNA isolated from lungs of SARS-CoV-2 infected hACE2, hACE2; *Irgm1*^−/−^ and hACE2; *Irgm1*^−/−^ + H-151 treated mice at 3 dpi. Mean ± SEM, ***p < 0.01, Brown-forsythe and Welch ANOVA (Dunnetts T3 multiple comparison test). **D.** Western blot analysis of NC protein expression in lung lysates of SARS-CoV-2 infected hACE2, hACE2; *Irgm1*^−/−^ and hACE2; *Irgm1*^−/−^ + H-151 treated mice at 3 dpi. **E-F.** qRT-PCR analysis for indicated genes with total RNA isolated from lungs of SARS-CoV-2 infected hACE2, hACE2; *Irgm1*^−/−^ and hACE2; *Irgm1*^−/−^ + H-151 treated mice (3 dpi). n=4, mean ± SEM ***p < 0.001, ****p < 0.0001, One-way ANOVA (Tukey’s multiple comparison test). **G.** Schematic representation of the experimental design depicting GSK583 treatment (25mg/kg body weight) followed by SARS-CoV-2 (5X10^4^ PFU) infection. Image created using Biorender.com. **H.** Graph showing total clinical scores of hACE2, hACE2; *Irgm1*^−/−^ and hACE2; *Irgm1*^−/−^ + GSK583 treated mice infected with SARS-CoV-2 for 5 days. Mean ± SEM, ****p < 0.0001, Two-way ANOVA (Tukey’s multiple comparison test). **I.** Viral copy number analysis in total RNA isolated from lungs of SARS-CoV-2 infected hACE2, hACE2; *Irgm1*^−/−^ and hACE2; *Irgm1*^−/−^ + GSK583 treated mice at 5 dpi. Mean ± SEM, *p < 0.05, **p < 0.01, Brown-forsythe and Welch ANOVA (Dunnetts T3 multiple comparison test). **J.** Western blot analysis of NC protein expression in lung lysates of SARS-CoV-2 infected hACE2, hACE2; *Irgm1*^−/−^ and hACE2; *Irgm1*^−/−^ + GSK583 treated mice at 5 dpi. **K-L.** qRT-PCR analysis for indicated ISGs with total RNA isolated from lungs of SARS-CoV-2 infected hACE2, hACE2; *Irgm1*^−/−^ and hACE2; *Irgm1*^−/−^ + GSK583 treated mice (5 dpi). n=4, mean ± SEM *p < 0.05, **p < 0.01, One-way ANOVA (Tukey’s multiple comparison test). **M.** Representative hematoxylin & eosin stained lung images of SARS-CoV-2 infected hACE2, hACE2; *Irgm1*^−/−^ and hACE2; *Irgm1*^−/−^ + GSK583 treated mice at 5 dpi.

**Supplementary Figure 9.**
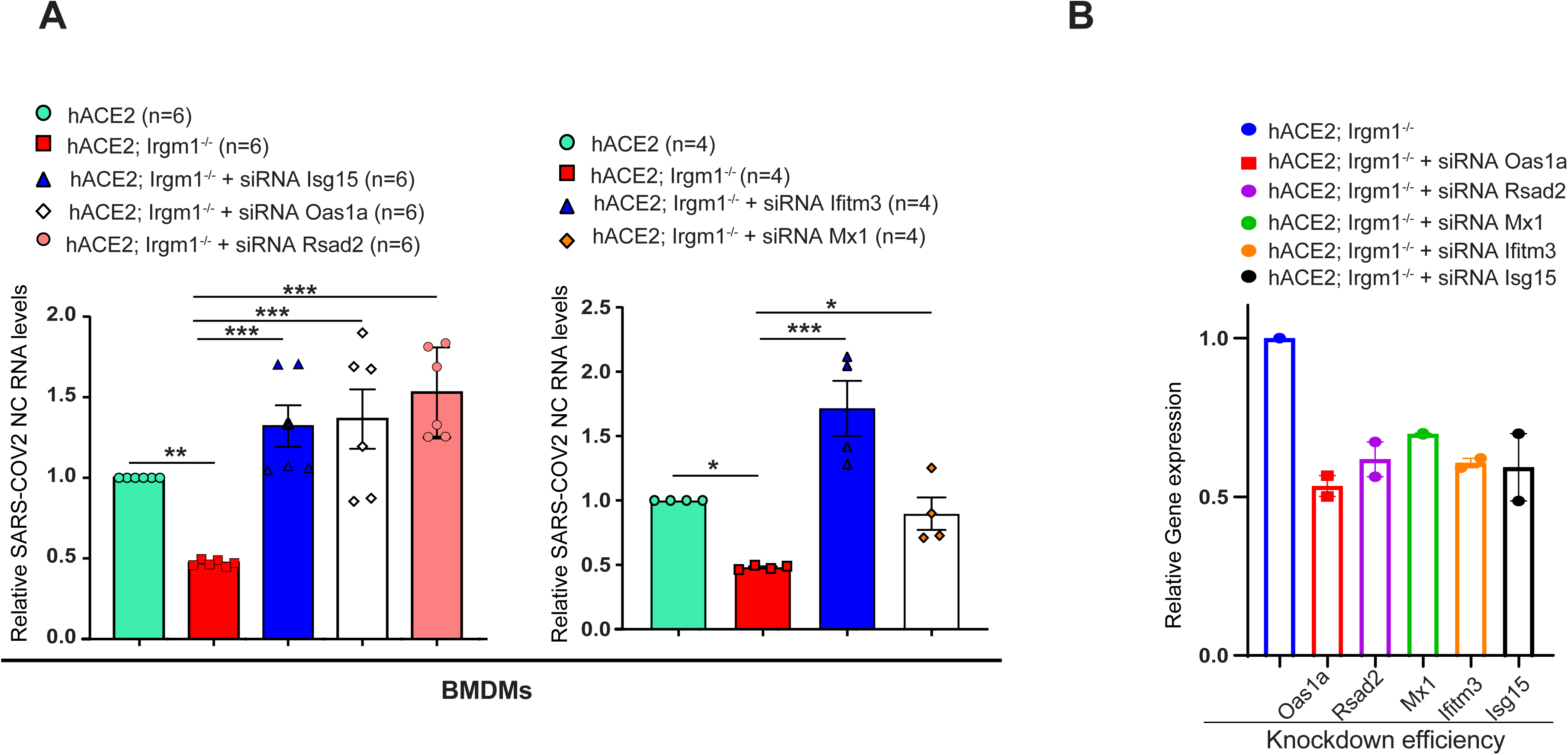
Depletion of anti-viral factors increases susceptibility of hACE2; *Irgm1*^−/−^ BMDM’s to SARS-CoV-2 infection. **A.** Total RNA was isolated from BMDMs of hACE2, hACE2; *Irgm1*^−/−^ transfected with control siRNA or Isg15 siRNA or Oas1a siRNA or Rsad2 siRNA or Ifitm3 siRNA or Mx1 siRNA as indicated followed by SARS-CoV-2 infection (MOI=0.5) for 12 hours and subjected to qRT–PCR with SARS-CoV-2 nucleocapsid specific primers (mean ± SEM, *p < 0.05, **p < 0.01, ***p < 0.001, Student’s unpaired t-test). **B.** Graph showing knockdown efficiency for each siRNA transfection in BMDMs.

## Supplementary Video Legends

**Supplementary Video 1.** The video of a representative SARS-CoV-2 infected hACE2 mouse on 6 dpi showing severe clinical symptoms such as constant body shaking, hunched posture, and labored breathing.

**Supplementary Video 2.** The video of a representative SARS-CoV-2 infected hACE2 and hACE2; *Irgm1*^−/−^ mouse on 6 dpi. Where hACE2; *Irgm1*^−/−^ were quite active with minor symptoms, the hACE2 mice had all severe symptoms and are not able to move.

**Supplementary Video 3** The video of a representative SARS-CoV-2 infected hACE2, hACE2; *Irgm1*^−/−^, hACE2; *Ifnar1*^−/−^ mouse on 6 dpi. Where hACE2; *Irgm1*^−/−^ mice were quite active with minor symptoms, the hACE2 mice had all severe symptoms and were not able to move. hACE2; *Ifnar1*^−/−^ were in the euthanasia stage with maximum clinical symptoms.

## Notes

### Competing Interest Statement

The authors have declared no competing interest.

